# Macrophage calcium signaling dynamics revealed using genetically-encoded sensors

**DOI:** 10.64898/2026.07.13.737685

**Authors:** Silke B. Chalmers, Teneale A. Stewart, Katherine Hughes, Anie Kurumlian, Mathilde Folacci, Krystyna A. Gieniec, Alexander J. Stevenson, Ngari Teakle, Anuj Sehgal, Emma Paydari, Jesper S. Thomsen, Katharine M. Irvine, Elvis Pandzic, Kate Poole, David A. Hume, Felicity M. Davis

**Author notes:** Correspondence to: David A. Hume and Felicity M. Davis. Equal contribution (Silke B. Chalmers, Teneale A. Stewart).

## Abstract

Macrophages (MΦs) inhabit all mammalian organs, adapting to and influencing the tissues in which they reside. These versatile cells rapidly respond to tissue-specific signals, clean up debris and serve as the first line of defense against pathogens. Their ability to act quickly relates to their abundance and specific arrangement in tissues, as well as the systems they possess for detecting, processing and integrating emergent signals. Ca^2+^ is a fast second messenger that has been linked to many of the core functions of MΦs. However, spatiotemporal features of physiologically-relevant Ca^2+^ signals in MΦs remain largely uncharacterized. Using mouse models expressing genetic biosensors and fluorescently tagged channels, we visualize MΦ Ca^2+^ signal dynamics and heterogeneity, uncover a remarkable degree of mechanosensitivity in these cells, and characterize the physiological consequences of genetic ablation of *Piezo1* channels via analysis of knockout models across their lifetime. Our in-depth investigation of MΦ Ca^2+^ signaling dynamics has broad relevance for the field of MΦ biology and the tissues that these cells support.

## Introduction

MΦs are an abundant resident cell population in every organ in the body (*1*). They are uniformly and regularly distributed and are associated with basement membranes of epithelia and endothelia, the surfaces of bone, muscle and connective tissue fibers, and the capsules of all major organs (*1*, *2*). Their survival, proliferation and differentiation are controlled by signals from the MΦ colony-stimulating factor receptor (CSF1R), a tyrosine kinase receptor expressed exclusively in the MΦ lineage (*3*). Transgenic reporter genes based upon regulatory elements of the *Csf1r* gene have enabled visualization of MΦs in many different tissue types (*4–9*). Resident MΦs are traditionally viewed as sentinels, providing the first line of defense against pathogens, but there is increasing evidence of their importance in homeostasis and sensing of tissue injury (*10*, *11*).

Regulated flux of Ca^2+^ across the plasma membrane and from intracellular Ca^2+^ storing organelles are important signal transduction pathways in nearly all mammalian cells (*12*). The emergence of cell-permeant, Ca^2+^-sensitive dyes in the 1980s (*13–15*) permitted visualization of global intracellular Ca^2+^ responses in immune cells and a subsequent eruption of studies describing ligand-mediated signal transduction events. One limitation to the use of these probes for the study of MΦ biology became evident very early, namely the relatively rapid extrusion of dyes from the cell by organic acid transporters (*16*). There have also been concerns that Ca^2+^-reporters dampen intracellular Ca^2+^ responses (*17*), and/or have off-target effects, potentially modifying or prolonging the response being measured (*18*). Notwithstanding these concerns, global changes in intracellular Ca^2+^ have been measured in MΦ and MΦ-like cell lines from mice and humans in response to a wide range of stimuli, including adhesion, migration, phagocytosis, and activation of purinergic and other plasma membrane receptors (reviewed in Ref (*19*)).

The ability of intracellular Ca^2+^ to regulate and coordinate diverse cellular processes throughout the body is attributable to the properties of this signal in space, time and magnitude (*20*). Ca^2+^ signals may arise due to release of Ca^2+^ from intracellular stores [e.g., by opening of inositol-1,4,5-trisphosphate receptors (InsP_3_Rs) on the endoplasmic reticulum (ER)] or by Ca^2+^ entry across the plasma membrane (e.g., via Ca^2+^ release-activated Ca^2+^ channels or ionotropic purinergic receptors). Store-operated Ca^2+^ entry (SOCE), involving ORAI channels and their classical activators, the ER Ca^2+^ sensors STIM1/2, has been detected in MΦs in response to a range of stimuli, including activation of endocytic and chemotactic receptors. However, knockout of STIM1/2 (producing complete inhibition of SOCE) demonstrate that this pathway is redundant in these cells (*21*). Ca^2+^ oscillations involving SOCE have also been implicated in the differentiation of osteoclasts (*22–24*). Regulated Ca^2+^ flux in MΦs is induced by numerous stimuli, including activation of purinergic receptors (e.g., P2RX7 and P2RX4) (*25*, *26*), C5AR1 (*27*), transient receptor potential vanilloid (TRPV) receptors 2 and 4 (*28*, *29*), and the channel-kinase TRPM7 (*30*, *31*), and is key to activation of the NLRP3 inflammasome (*32*, *33*). More recently, the mechanosensitive, non-selective cation channel PIEZO1 was added to the set of Ca^2+^-permeable regulatory proteins in MΦs during various pathologies (*34*, *35*).

An alternative to the use of Ca^2+^-sensitive dyes for the visualization of intracellular Ca^2+^ came with the introduction of genetically-encoded Ca^2+^ indicators, e.g., GCaMP and its derivatives, which are based upon fusions of calmodulin, the calmodulin binding M13 peptide and green fluorescent protein (*36–38*). The GCaMP scaffold has since been modified by numerous groups, to optimize speed, sensitivity and brightness (*39*). GCaMP6f emerged to enable ultrasensitive imaging of rapid Ca^2+^ transients associated with neuronal activity (*40*). The majority of studies utilizing GCaMP6f to date have focused on Ca^2+^ signals associated with neuronal activity *in vitro* and *in vivo*. However, important studies in astrocytes, cardiomyocytes, and epithelial cells have illustrated its potential for imaging sub-cellular Ca^2+^ events, as well as fast Ca^2+^ transients and large intercellular responses, across a range of cell and tissue types (*41–44*). We reasoned that expression of GCaMP6f in MΦs could offer a valuable platform for high resolution visualization and analysis of regulated Ca^2+^ flux *in situ* and *in vitro* in response to a range of physiologically relevant stimuli. For this purpose, we took advantage of a Cre-dependent GCaMP6f line (*45*) and a *Csf1r*-cre line (*46*) that was anticipated to drive reporter expression in cells of the MΦ lineage. This model enabled us to visualize Ca^2+^ events from the scale of sub-cellular dynamics to the level of tissue-wide organization.

## Results

### Spatiotemporal dynamics of Ca^2+^ responses in tissue MΦs

The *Csf1r* promoter and enhancer have been used extensively to drive reporter genes (*4*, *47*) as well as a tamoxifen-inducible cre transgene (*48*). We wished to avoid the use of tamoxifen, which can have direct effects on MΦ function (*3*, *49*). The specificity of expression of cre in the *Csf1r-icre* line (*50*) was originally determined by crossing to a conditional *Rosa26-YFP* reporter line. Yellow fluorescent protein (YFP) was detected in various tissues including liver, spleen, intestine, heart, kidney, and muscle. Based upon localization of YFP in a subset of these tissues, the signal was inferred to be MΦ-associated. We generated mice expressing *Csf1r-cre* in combination with a Cre-dependent Ca^2+^ sensor *GCaMP6f* and Ca^2+^-independent red fluorescent protein *tdTomato* (*tdT*) at the Rosa26 locus (herein referred to as *GCaMP6f^+^;tdT^+^;Csf1r-cre^cre/+^*mice) (**Fig 1A-B and S1A**). We observed idiosyncratic expression of tdT-positive cells in some tissues (**Fig S1B**), most notably the pancreas and peri pancreatic fat, which had a visibly pink hue (**Fig 1B-C**), that had not been reported in prior studies (*51*). This finding raises questions regarding the interpretation of an earlier study investigating the role of STAT3 in tumorigenesis (*50*) and a subsequent study that used *Csf1r-icre* to claim that *Csf1r* can be expressed in neurons (*52*). Nevertheless, the expression of tdT in regularly spaced stellate cells in the skin, salivary gland and mammary gland (**Fig 1B**) resembled the location of *Csf1r* reporter transgenes, including using a knock-in CSF1R-FusionRed transgene that reports protein expression (*53*).

**Fig 1:**
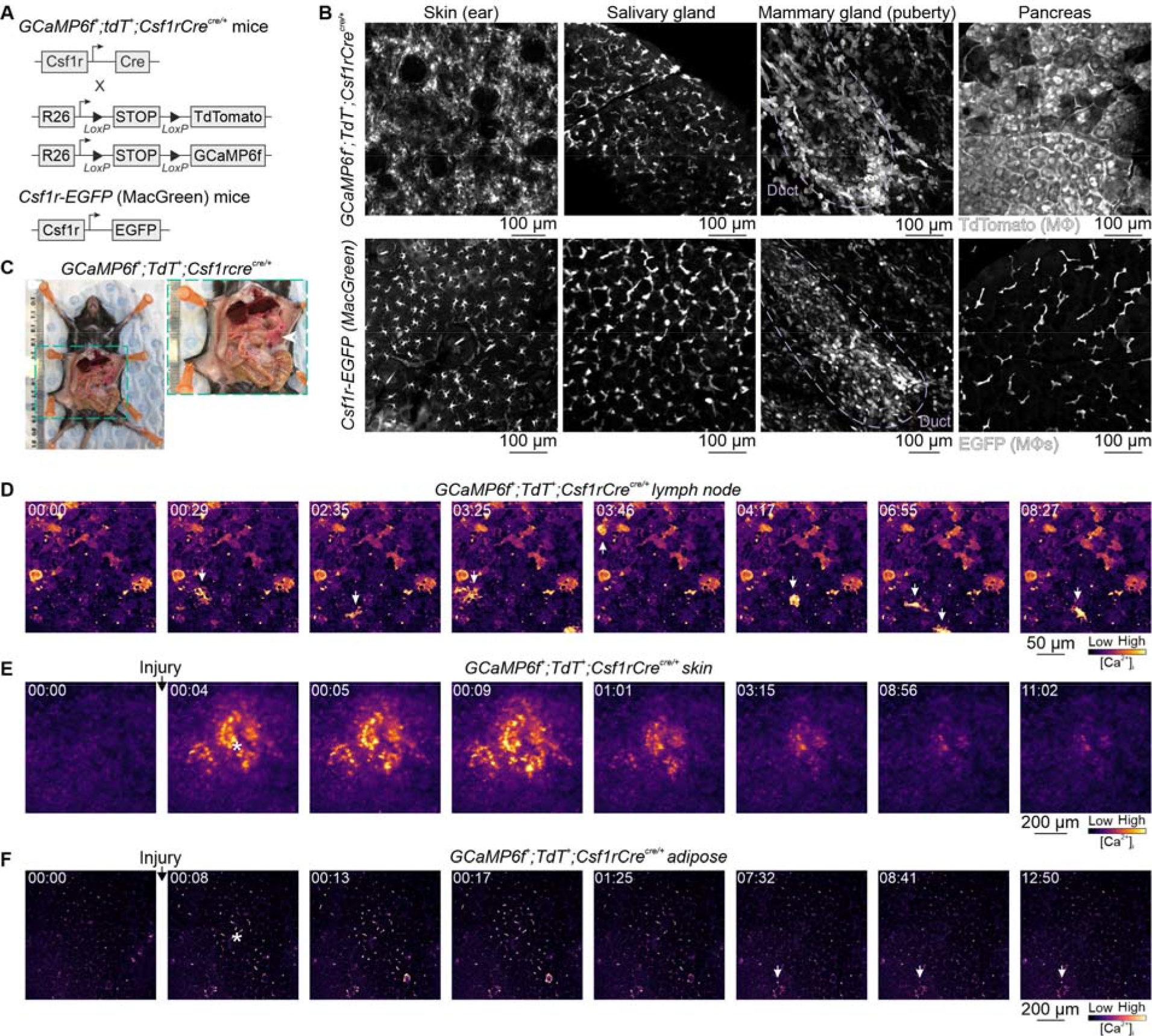
*GCaMP6f^+^;tdT^+^;Csf1r-cre^cre/+^*model development and characterization. **A**, The strategy for generating *GCaMP6f^+^;tdT^+^;Csf1r-cre^cre/+^* mice (see also Fig S1). **B**, A comparison of *GCaMP6f^+^;tdT^+^;Csf1r-cre^cre/+^* and *Csf1r-EGFP* (MacGreen) models. Images are maximum intensity projections of individual optical slices showing tdT^+^ cells (top panel) or EGFP^+^ cells (bottom panel) in the skin, salivary gland, pubertal mammary gland and pancreas (see also Fig S1). **C**, Image showing the abnormal pinkish hue of pancreas, and peri pancreatic fat, from *GCaMP6f^+^;tdT^+^;Csf1r-cre^cre/+^*mice. Live imaging of spontaneous intracellular Ca^2+^ activity in lymph nodes (**D**) and mechanically-induced intracellular Ca^2+^ recordings in ear skin (**E**) and adipose tissue (**F**). Arrows in D highlight spontaneously active cells. Asterisk in E and F shows point of mechanical stimulation; [min:sec]; *n* = 3–4.

In mice, GCaMP6 has been used in microglia to visualize intracellular Ca^2+^ responses to acute neuronal activation *in vivo* and purinergic stimulation in brain slices (*54*, *55*). It has also been used to characterize Ca^2+^ dynamics of meningeal MΦs, and GCaMP5 has been used to visualize Ca^2+^ signals in *Csfr1-Cre* positive cells positioned around injected tumour cells (*51*). We first determined whether GCaMP6f was sufficient to detect Ca^2+^ responses in other whole tissue preparations. Spontaneous Ca^2+^ activity was observed in GCaMP6f expressing cells in lymph nodes (**Fig 1D** and **Movie S1**). To mimic a mechanical stimulus, we lightly touched different tissue types with the tip of a needle. In both skin and adipose tissue, we observed fast Ca^2+^ signals in GCaMP6f-positive cells, notably in epidermal Langerhans cells, where stimuli of this intensity would not be uncommon (**Fig 1E-F and Movies S2-3**). Ca^2+^ signals ramified away from the point of contact and were transient in nature, suggesting a sensitive response to a minimally-invasive force, likely involving biochemical and mechanical aspects.

### MΦ *single cell Ca^2+^ responses*

We next isolated peritoneal MΦs from *GCaMP6f^+^;tdT^+^;Csf1r-cre^cre/+^* mice and generated bone marrow-derived macrophages (BMDMΦs) from femoral bone marrow. Expression of tdT and GCaMP6f fluorescent proteins were confirmed in F4/80-positive BMDMΦs using endogenous tdT fluorescence (**Fig 2A**) and an anti-GFP antibody recognizing GCaMP6f (*56*) (**Fig S2A**). Phagocytosis by immune cells is believed to be associated with local elevations in intracellular Ca^2+^ (*57*). However, to our knowledge, this has never been confirmed in tissue MΦs with Ca^2+^ imaging with subcellular resolution. GCaMP6f was sufficiently sensitive to detect localized Ca^2+^ events associated with particle uptake in MΦs (**Fig 2B**, blue box) but limitations on temporal resolution under these imaging conditions precluded a detailed characterization of the nature of wave propagation from the phagocytic cup. Nevertheless, we observed that peritoneal MΦs exposed to fluorescent zymosan exhibited large cytosolic Ca^2+^ oscillations as they interacted with and ultimately engulfed fluorescent particles (**Fig 2B** and **Movie S4**). Ca^2+^ oscillations continued for approximately 40 s and resumed when MΦs engulfed subsequent units.

**Fig 2:**
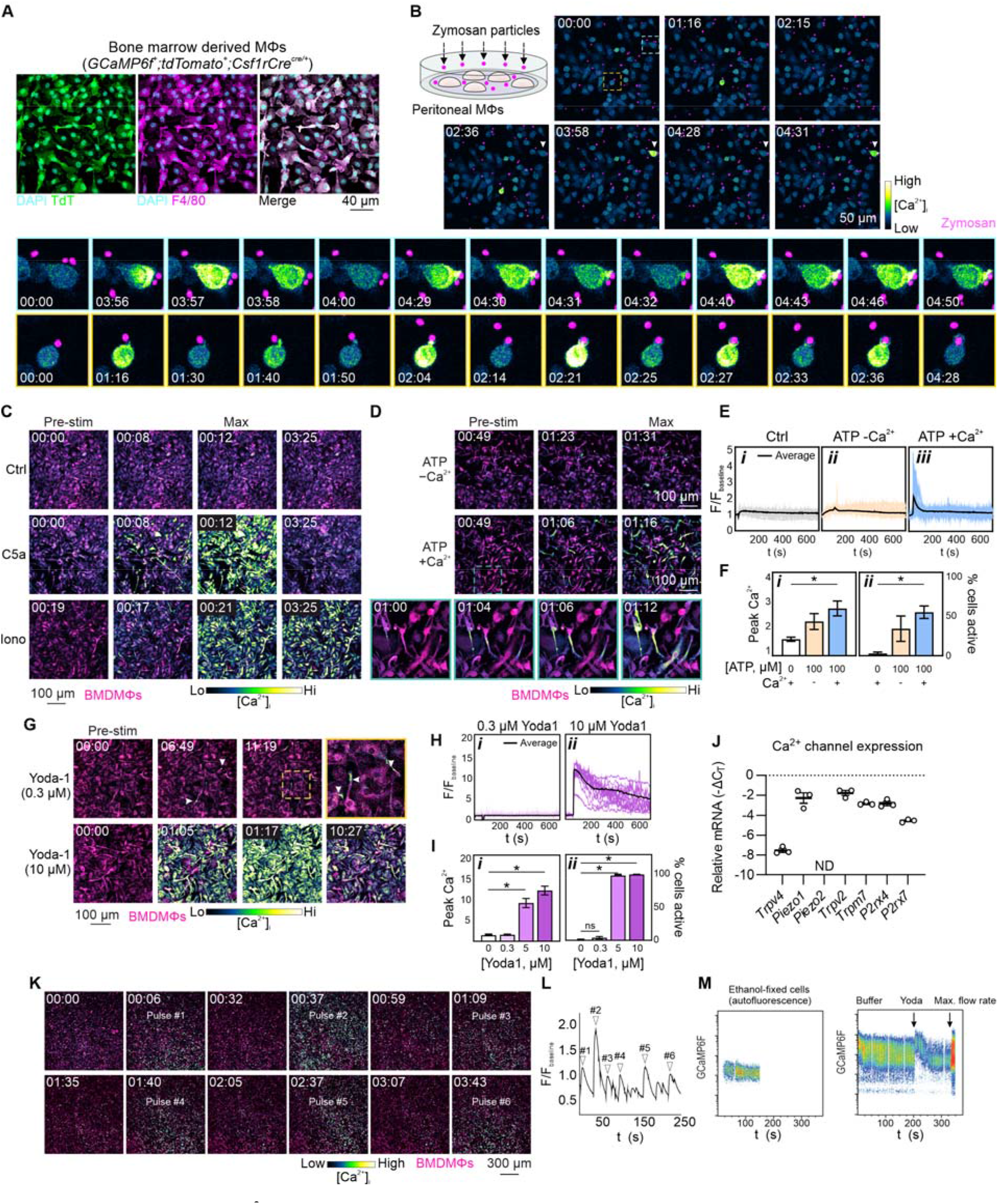
Single cell Ca^2+^ signaling in MΦs. **A**, tdT^+^ (green) bone marrow derived MΦs immunostained with anti-F4/80 antibody (magenta); nuclei are stained with DAPI (cyan). **B**, Intracellular Ca^2+^ responses in peritoneal MΦs exposed to fluorescent zymosan (magenta). Intracellular Ca^2+^ responses in BMDMΦs stimulated with (**C**) buffer only control, C5a (100 nM) and ionomycin (iono, 10 μM) and (**D**) ATP (100 μM ± extracellular Ca^2+^). Single cell Ca^2+^ traces (average in black) in response to ATP (100 μM ± extracellular Ca^2+^) (**E**) and quantification showing peak Ca^2+^ response and percent responding cells (**F**). **G**, Intracellular Ca^2+^ responses in BMDMΦs stimulated with Yoda-1 (0.3 and 10 μM), quantified in **H-I**. **J**, BMDMΦ mRNA levels (-ΔC_T_) of ion channels implicated in MΦ biology. ND, not detected. **K**, Intracellular Ca^2+^ responses in BMDMΦs following repeated flushing with physiological salt solution. **L** Population average Ca^2+^ responses to fluid flow pulses shown in **K. M,** GCaMPP6f signal was measured during flow cytometry acquisition of ethanol fixed BMDMΦs (first panel) or live cells (second panel). Live BMDMΦs were treated with Yoda-1 at 200 s acquisition, and the flow rate was increased from 10 µL/minute to 60 µL/min at 300 s. Bar graphs show mean ± s.e.m, *n* = 3 independent experiments; [min:sec]. Significance was assessed via ordinary one-way ANOVA with Tukey’s multiple comparison test, \**p* < 0.05.

We next explored MΦ responses to agents that have previously been linked to Ca^2+^ signaling in primary MΦs and/or cell lines. C5a, a central effector produced through complement activation, binds to the G-protein coupled receptors, C5aR1 and 2. Consistent with previous findings in fura2-loaded MΦs (*58*), C5a (100 nM) caused a large transient increase in intracellular Ca^2+^ in nearly all GCaMP6f-expressing BMDMΦs (**Fig 2C, S2B-C, Movie S5-6**). The Ca^2+^ response was similar in nature (peak, % responding) to the Ca^2+^ response caused by the Ca^2+^ ionophore ionomycin, which was sustained (**Fig 2C**, **S2B-C and Movie S7**). By comparison, relatively high concentrations of the purinergic receptor agonist adenosine triphosphate (ATP), known to activate ionotropic P2X and metabotropic P2Y receptors in MΦs (*59*), produced Ca^2+^ responses in only ∼50% of cells bathed in Ca^2+^-containing buffer (**Fig 2D-F**, **Movie S8**), suggestive of intriguing heterogeneity in purinergic receptor expression, modulation and/or localization amongst neighboring BMDMΦs. The ATP-induced Ca^2+^ signal propagated through these cells as a slow (6–8 s) intracellular wave (**Fig 2D**).

The ion channel PIEZO1, identified by Coste et al. in 2010 (*60*), has since been linked to numerous processes involved in the MΦ response to infection or injury, including bactericidal activity (*61*), polarization (*35*), efferocytosis (*62*) and phagocytosis (*63*). This mechanosensitive Ca^2+^-permeable ion channel is activated by the synthetic small molecule Yoda-1. Consistent with this, GCaMP6f-expressing BMDMΦs showed large, measurable cytosolic responses to 5 and 10 μM (but not 0.3 μM) Yoda-1 (**Fig 2G-I**). Robust Yoda-1 responses were also observed in tissue resident peritoneal MΦs (**Fig S2D**). Intracellular Ca^2+^ levels remained elevated for several minutes after Yoda-1 stimulation and exhibited decaying oscillations (**Fig 2H** and **Movie S9**). *Piezo1*, but not *Piezo2*, mRNA was detectable in BMDMΦs, as were other Ca^2+^-permeable channels widely studied in this cell type (**Fig 2J**). This prompted us to further explore the signaling and function of PIEZO1 in primary MΦs and its potential regulation of MΦ function *in vivo* under physiological conditions.

### *M*Φ Ca^2+^ responses to fluid shear stress

PIEZO1 is involved in the sensing of shear stress in endothelial cells and shear stress-mediated elevations in cytosolic Ca^2+^ (*64*, *65*), which are phenomena important for normal vascular development and homeostasis (*66*, *67*). To better understand the mechanisms by which MΦs sense shear stress, we took the simple qualitative approach of vigorously flushing an adherent monolayer of MΦs with physiological salt solution and recorded resulting intracellular Ca^2+^ responses (**Fig 2K, Fig S2E** and **Movie S10**). Using this approach, we detected robust intracellular Ca^2+^ responses to fluid shear stress in adherent BMDMΦs (**Fig 2K-L**). To assess whether shear stress also increases in Ca^2+^ in MΦs in suspension, we investigated whether a similar response could be detected using flow cytometry. Indeed, GCaMP6f-expressing MΦs produced a fluorescent signal that was dependent on flow rate (**Fig 2M**).

### Large PIEZO1-mediated currents in isolated MΦ*s*

Ionic currents can be mechanically evoked in cells *in vivo* in response to a range of physiological stimuli, including shear stress, stretch and compression, as well as by cell-generated forces, which can be modelled and studied in various assays (*68*)). Previous studies have demonstrated that small, high threshold PIEZO1-mediated currents can be evoked by the application of membrane stretch to BMDMΦs (*69*). Here, we evoked stretch-activated currents with a similar kinetic profile and appearance to these reports (**Fig 3A**). However, the kinetics and sensitivity of these currents were distinct from those reported for PIEZO1 activation in many other cells and tissues, potentially relating to differences in PIEZO1 interacting partners among these different cell types and subtypes (*60*, *70*, *71*) (**Fig 3B-C**). When stimuli were applied by displacing substrate elements subadjacent to the cell (*72*, *73*), however, sensitive currents were evoked (**Fig 3D-E**). As may be expected of this cell type, BMDMΦs plated on these arrays would occasionally try to phagocytose individual pili, and caution had to be exercised (using the tdT signal) to avoid compressing these regions. Further, when the stimulus produced a current that did not return to baseline or evoked a current with stimuli adjacent to the pilus (in the absence of pilar movement), the stimulation point was discarded (9 out of 23 cells). In all remaining cells (14 of 14), currents were evoked by substrate deflections. Current kinetics (latency, activation time constant, inactivation time constant) were consistent with previous measurements of PIEZO1 activity using pillar arrays (**Fig S3**) (*71*, *73*, *74*), and the latency and activation time constant were suggestive of a direct coupling between mechanical input and channel activation. The median threshold of activation (calculated as the smallest applied stimulus that evoked a current) for these currents was measured at 24.8 nm (mean ± s.e.m. = 31 ± 8 nm).

**Fig 3:**
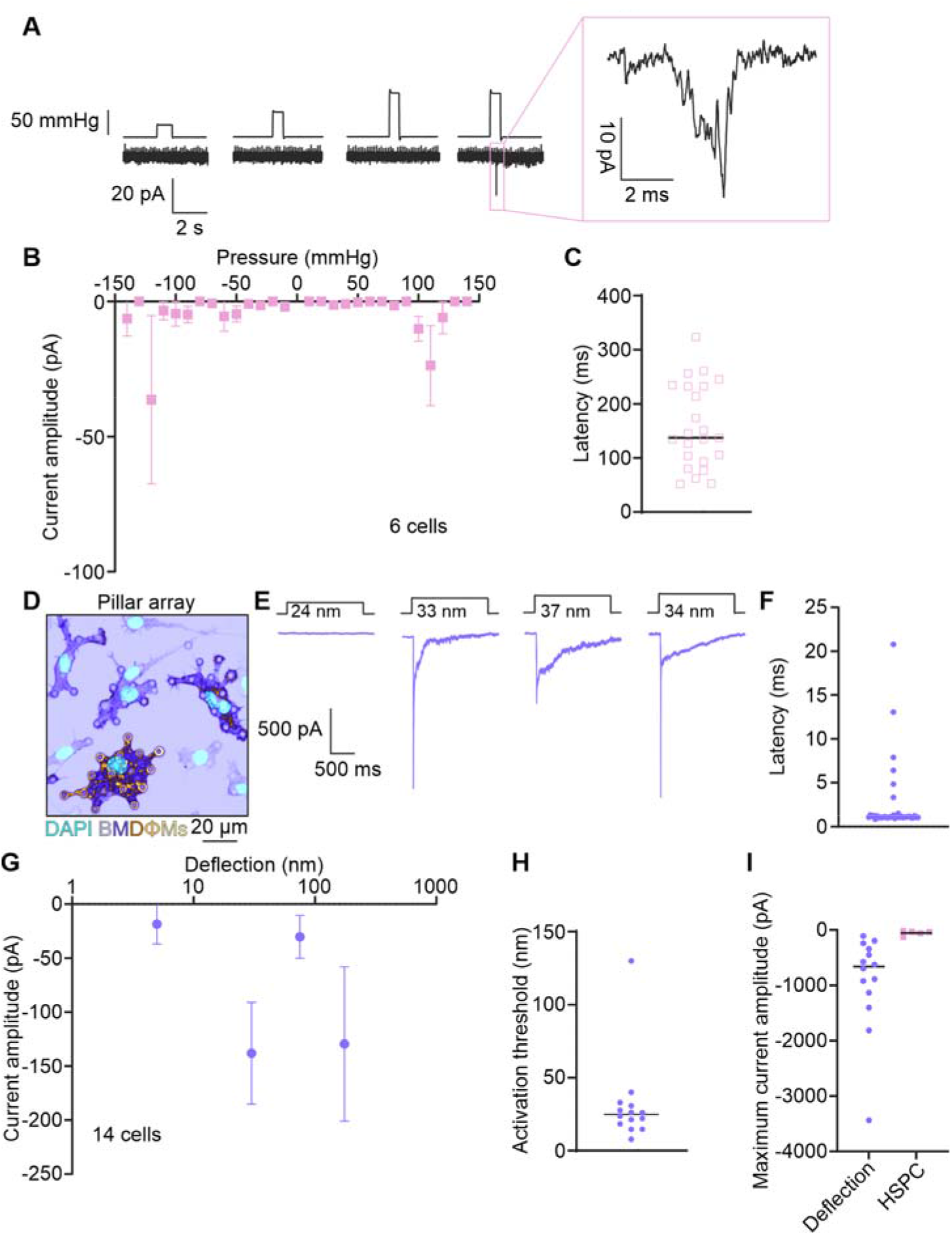
Mechanically-evoked currents in BMDMΦs. **A**, Representative traces of mechanically-evoked currents in BMDMΦs from an outside-out patch in response to increased positive pressure from 25 to 95 mmHg, example trace expanded in insert (pink box). **B**, Stimulus-response plot for BMDMΦs cultured on pillar arrays recorded in outside-out patches using high speed pressure clamp (HSPC) (*n* = 6 cells, data is presented as mean ± s.e.m). **C**, Latency of currents evoked using HSPC (*n* = 23 currents, median = 137.3 ms). **D**, Representative image of BMDMΦs grown on elastomeric pillar array and fixed and stained with DAPI (nucleus) and F4/80 (BMDMΦs). **E**, Representative traces of mechanically evoked currents in BMDMΦs cultured on pillar arrays in response to serial deflections. **F**, Latency of currents evoked by pillar deflections (*n* = 37 currents, median =1.090 ms). **G**, Stimulus-response curve for BMDMΦs cultured on pillar arrays in response to pillar deflections (*n* = 14 cells, data is presented as mean ± s.e.m). **H**, Activation threshold of deflection evoked currents (*n* = 14 cells, median = 24.80 nm). **I**, Maximum current amplitude in each cell in response to mechanical stimulus. (purple circles = pillar deflection, *n* = 14 cells, median = −662.1 pA; pink squares = HSPC, *n* = 5 cells, median=−38.80 pA).

These data confirm that mechanically evoked currents, activated within the cell-substrate interface in BMDMΦs, exhibit the hallmarks of direct activation by mechanical input and are characterized by a low threshold for activation (**Fig S4**) (*35*). Such exquisite sensitivity of mechanically evoked currents has previously been measured in low threshold somatosensory neurons (*73*), in additional systems where PIEZO1 is expressed alongside accessory molecules that sensitize the channel (i.e. STOML3) (*73*) and when heterologously expressed in cells with high prestress (*74*).

### Histopathological assessment of young and aged mice with Piezo1 knocked out in CSF1R^+^ MΦs

The expression of *Piezo1* in BMDMΦs, the sensitivity of these cells and tissue resident MΦs to Yoda-1, and the remarkable mechanical activation of BMDMΦs at the cell-substrate interface, prompted us to investigate *in vivo* roles for this channel in tissue MΦs under non-perturbed (physiological) conditions. We generated mice with conditional knockdown of *Piezo1* in *Csf1r*-expressing cells (*Piezo1^fl/fl^;Csf1rCre^Cre/+^*or “cKO” mice) and confirmed knockdown at the mRNA level (**Fig 4A**). No changes were detected in mRNA levels of other MΦs enriched targets (*Trpv4* and *Trpv2*), *Piezo2* (which remained undetected) or other Ca^2+^ channels and binding proteins that were assessed. We stimulated BMDMΦs from control and cKO mice with Yoda-1 (10 μM), confirming knockdown of functional PIEZO1 channels in this model (**Fig 4B-D**).

**Fig 4:**
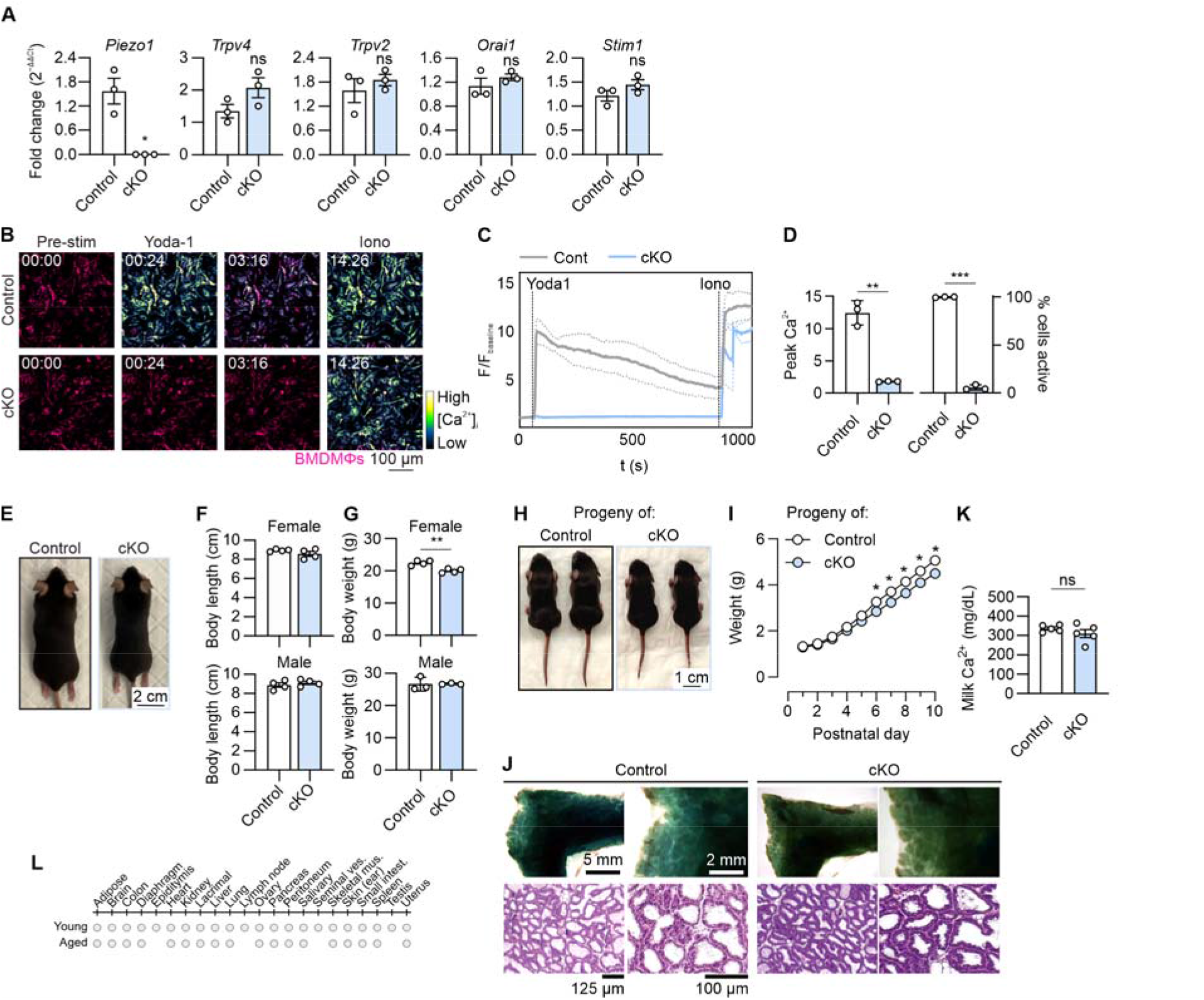
Characterization of *Piezo1^fl/fl^;Csf1r-cre^Cre/+^*model under non-stressed conditions. **A**, BMDMΦ mRNA levels of *Piezo1* and other Ca^2+^ signaling associated proteins in cKO and control animals. **B,** Intracellular Ca^2+^ responses to 10 µM Yoda-1 and subsequent 10 µM ionomycin addition in BMDMΦs isolated from control and cKO animals, quantified in **C–D**. **E**, Representative visual comparison of control and cKO animals, with body length and weight measurements compared in **F** and **G**. **H**, Representative comparison of pups (without *Piezo1* deletion in MΦs) nursed by control or cKO dams, with pup weight quantified in **I**. **J**, Mammary gland wholemount and histology from lactating control and cKO dams. **K**, Comparison of milk Ca^2+^ concentration in samples obtained from control or cKO nursing dams. **L**, Summary of tissue histology assessed by veterinary pathologist from young (12–14 weeks) or aged (52 weeks) control and cKO animals. Grey circles indicates organs that were assessed and determined to have no major differences. Graphs are mean ± s.e.m, n = 3 (**A–D**), 4 (**F**), 5 (**J, K, L**), or 6 (**I**) independent experiments. Significance was assessed via unpaired t-test (**D, F, G, K**), or two-way ANOVA with Bonferroni post-test (**I**), \**p* < 0.05. [min:sec].

No marked difference in the outward appearance (**Fig 4E**), body length (**Fig 4F**), or femoral length (**Fig S5A-B**) were observed in cKO mice compared to control. A small but significant difference was observed in the body weight of adult female, but not male, cKO mice (**Fig 4G**) as well as in pups with normal *Piezo1* expression that were nursed by cKO female mice (**Fig 4H-I**). CSF1R-dependent MΦs are required for normal mammary development and maturation (*75*). However, the reduced weight of post-pubertal female mice did not correspond with a change in mammary histology or wholemounts during lactation (**Fig 4J**) or in milk Ca^2+^ levels (**Fig 4K**). While PIEZO1 signaling has been implicated in the regulation of osteoclast differentiation (*76*) and bone formation (*77*), and female mice of this background have been shown to develop early onset osteoporosis (*78*), our microtomographical bone measurements from post-lactational females revealed no meaningful impact of *Piezo1* deletion (**Fig S5A, C-G**).

A comprehensive, blinded histological analysis of tissue from young and aged mice, performed by a veterinary pathologist, revealed no marked changes in >20 organs assessed (**Fig 4L**), and examination of selected tissues with IBA1 immunostaining showed no difference in MΦs distribution (**Fig S6**). Finally, we observed no statistical differences in the early or late uptake of zymosan particles in peritoneal MΦs (percent positive or particle number, **Fig S7**). Thus, zymosan-evoked [Ca^2+^]_i_ responses in MΦs are not caused by Ca^2+^ release-activated channels (shown by Vaeth et al., in Ref (*21*)) or by PIEZO channels (**Fig S7**). Collectively, these data suggest that PIEZO1 function in MΦs is dispensible for trophic or homeostatic functions *in vivo*.

### Dynamic properties of PIEZO1 channels in wounded MΦ*s*

MΦs are an abundant cell type in all tissues and are recruited during injury and infection (*1*, *2*). Our electrophysiological recordings support roles for PIEZO1 at the MΦ cell-substrate interface, leading us to investigate whether PIEZO1 regulates MΦ movements during injury-mediated recruitment. Our investigations in scratch-wounded BMDMΦs revealed three key findings. Firstly, we observed a robust increase in intracellular Ca^2+^ in cells on the wound edge in response to first and subsequent injuries (**Fig 5A and Movie S11**). The amplitude and temporal profile of the response was shaped by the presence of extracellular Ca^2+^ (**Fig 5B and Movie S12**). However, Ca^2+^ events were still observed in cells bathed in Ca^2+^-free buffer (**Fig S8**), suggesting a role for Ca^2+^ mobilization from intracellular stores. While locally released ATP is a likely candidate in this damage response (*79*), through both ionotropic and metabotropic purinergic receptors, all cells on the wound edge respond with an increase in Ca^2+^, in contrast with Ca^2+^ recordings in ATP-stimulated BMDMΦs (**Fig 2D**), implying additional mechanisms of activation. Secondly, we made the interesting observation that after BMDMΦs enter the denuded area, these cells often oriented themselves in a “patrol line” along the wound edge (**Fig 5C and Movie S13**). By examining channel movements in patrolling BMDMΦs isolated from *Piezo1-tdT* mice (expressing tdT fused to the C-terminus of PIEZO1 (*66*)), we observed PIEZO1 channels to transiently coalesce at sites where BMDMΦs meet (**Fig 5D and Movie S14**). We occasionally observed PIEZO1 channels in the trailing edge of migrating BMDMΦs (**Fig S9A** and **Movie S15**), similar to that observed in migrating keratinocytes (*80*), but we rarely observed PIEZO1 channels to accumulate along inert surfaces, such as silicone dish inserts (**Fig S9B** and **Movie S16**). These results demonstrate that, in adherent motile MΦs, PIEZO1 channels rapidly and transiently assemble at sites of dynamic cellular encounters.

**Fig 5:**
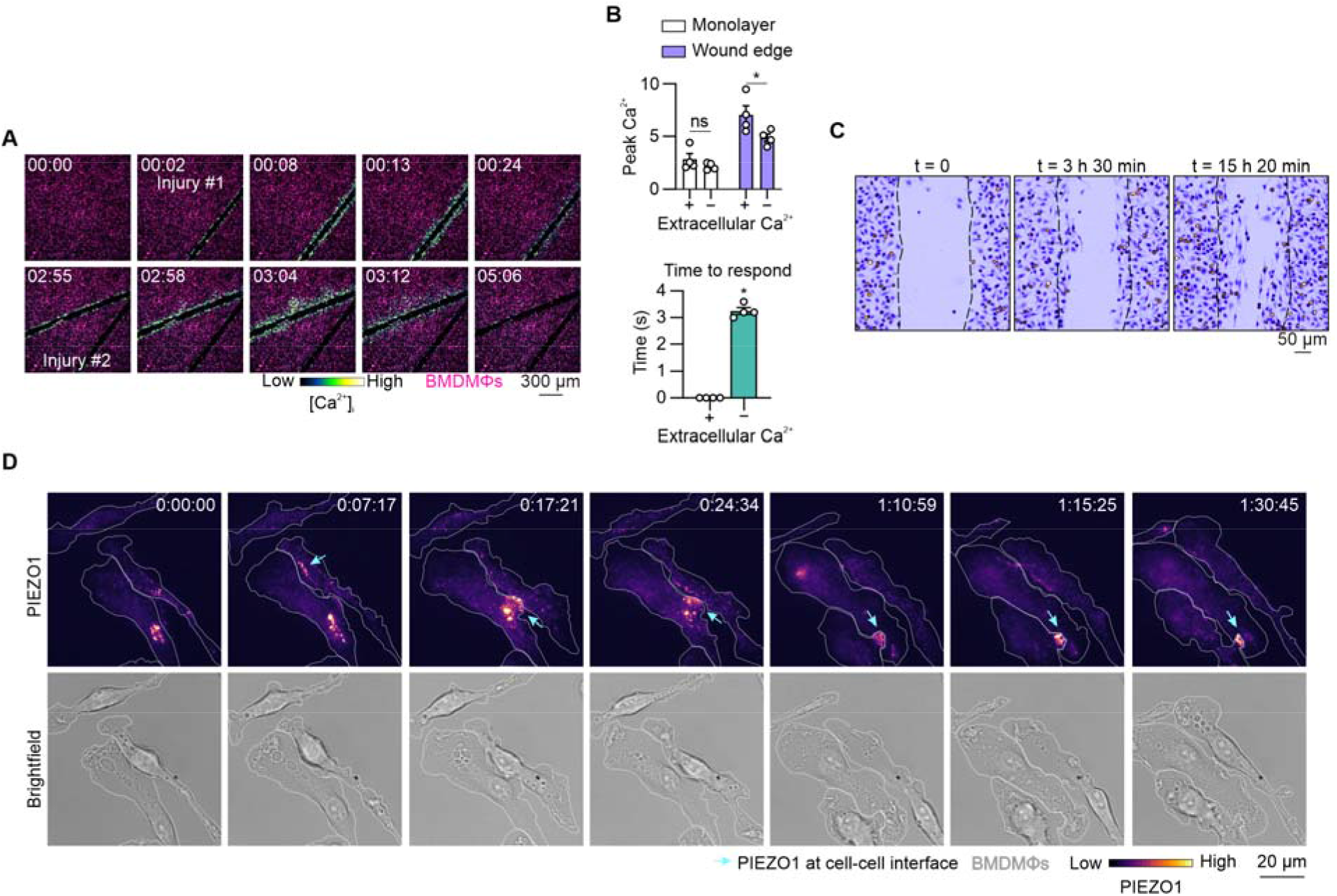
BMDMΦ dynamics in wounding and cell contacts. **A**, Ca^2+^ responses at the wound edge of BMDMΦ monolayers bathed in 1.8 mM extracellular Ca^2+^, quantified in (**B**) ± Ca^2+^. **C**, BMDMΦ migratory behavior along monolayer wounds. **D**, PIEZO1 channels coalesce at sites of BMDMΦ cell-cell contacts. Bars are mean ± s.e.m., *n* = 3 independent experiments. Significance assessed via unpaired t-test, \**p* < 0.05. [h:min:sec].

## Discussion

Our generation of transgenic mice expressing the sensitive genetically-encoded Ca^2+^ indicator GCaMP6f in MΦs has provided an opportunity to study chemically- and mechanically-evoked Ca^2+^ signaling events in these cells at high spatial and temporal resolution *in vitro* and *in situ*. A previous study has used the *Cx3cr1-Cre* driver to express GCaMP6s or a membrane tethered version of GCaMP6f in microglia (*55*). These authors detected spontaneous, localized Ca^2+^ transients and increased signaling in response to changes in neuronal activity. We used *Csf1r-Cre* to express a fast Ca^2+^ reporter for visualization of MΦs in the periphery. The *Csf1r-Cre* proved less lineage-specific than expected (*46*, *51*). Nevertheless, using this model in select cells and tissues, we were able to detect spontaneous Ca^2+^ events in *ex vivo* preparations and elicit a propagated response to mechanical stimulation. Our findings suggest that mechanosensing is a common property of tissue resident MΦs and pave the way for future studies using implantable imaging windows, such as those employed by Taghdiri et al. (*51*), to observe and monitor acute-and long-term MΦ responses to a range of physiological and pathological stimuli. These experiments necessitate careful implantation of imaging windows to maintain tissue sterility and avoid MΦ activation prior to imaging (*79*), and may benefit from the use of flexible or openable window technologies for chemical and/or mechanical stimulation (*81*, *82*).

Our *in vitro* investigations demonstrated that the GCaMP6f reporter was sufficiently sensitive to detect intracellular Ca^2+^ waves within individual MΦs. We were able to localize Ca^2+^ transients to the vicinity of the phagocytic cup associated with particle uptake in peritoneal MΦs and observe decaying oscillations following stimulation with the PIEZO1 agonist Yoda1 in BMDMΦs. Interestingly, we detected Ca^2+^ responses arising from experimental manipulations that are routinely carried out in the study of MΦ biology – flushing of cells on the culture plate and flow cytometry. However, it is not yet clear whether such manipulations lead to temporary or sustained alterations in cellular function nor is it clear what the consequences of these perturbations are on downstream cellular readouts. In addition, we observed propagating intracellular Ca^2+^ signals in response to supraphysiological concentrations of extracellular ATP. We uncovered that not all cells utilize intracellular Ca^2+^ to respond to ATP, demonstrating an intriguing degree of signaling heterogeneity that may be a cause or consequence of MΦ functional diversity (*83*). Notably, robust Ca^2+^ signals were consistently observed in cells adjacent to the wounded edge of cell monolayers, where ATP is released locally in response to cell damage (*84*, *85*). However, in this case, all cells in the immediate vicinity exhibited large intracellular Ca^2+^ responses, suggesting that additional (purinergic receptor-independent) factors are involved in the MΦs response to wounding.

We focused on mechanically evoked ion fluxes and the basic biology of PIEZO1 in MΦs. In contrast to the currents produced by stretching BMDMΦs at the apical surface, we observed exceptional sensitivity of mechanically evoked currents at the cell-substrate interface. Such striking sensitivity to mechanical inputs is typically only seen in low threshold somatosensory neurons or in systems where PIEZO1 is expressed alongside accessory molecules that sensitize the channel (*73*, *86*). Whilst our study focused on PIEZO1 channels in MΦs, these cells also express TRPV4, and signaling at the cell-substrate interface may reflect the combined activity of these two channels, as previously reported in primary chondrocytes (*71*). Nevertheless, taken together, our results suggest that BMDMΦs have the capacity to distinguish between distinct types of physiologically-relevant mechanical stimuli, which may be relevant as these highly motile cells infiltrate diverse tissue environments.

One study published in 2024, which also examined PIEZO1 activation in BMDMΦs, reported that the Yoda1-mediated intracellular Ca^2+^ signals observed in these cells were not present in tissue resident (cardiac) MΦs (*28*). Based on this observation, the authors concluded that BMDMΦs fundamentally differ from tissue resident MΦs, suggesting that the differences observed between these two cell types could be an artifact of the *in vitro* differentiation process. In contrast, we observed robust Yoda1 responses in both BMDMΦs and freshly isolated peritoneal MΦs, suggesting that tissue MΦs can indeed exhibit cellular responses to Yoda1. We suggest that the differences observed between BMDMΦs and cardiac MΦs (isolated by enzymatic digestion and purified by flow cytometry) may reflect the chemical and mechanical stress imposed by these processes, or alternatively the markedly different cellular environments experienced by distinct tissue resident MΦs populations.

Recently, PIEZO1 has been shown to have numerous roles in the MΦ mediated defense against disease or infection (*34*, *61*, *62*). However, to our knowledge, no study has comprehensively characterized potential roles for this channel in these cells during tissue development and homeostasis. MΦs offer a fascinating and relevant system to study PIEZO1 function, as they are highly motile cells exposed to a range of tissue properties, and exhibit diverse phenotypes and distribution patterns (*1*). To determine the homeostatic function of PIEZO1 in tissue MΦs, we generated conditional knockout mice and studied them across their lifetime. Effective loss of function was demonstrated by the complete loss of response to Yoda1, complementing our gene expression analysis. However, we were not able to detect biologically relevant changes in MΦ distribution or function in the absence of PIEZO1, including in bone, where one may anticipate an important role in osteoclasts (*87*). Moreover, intracellular Ca^2+^ responses were detected in tissue resident peritoneal MΦs derived from control and cKO mice and no change in fluorescent particle uptake was observed in our study. This is contrast to a modest reduction in particle uptake by BMDMΦs isolated from loss-of-function *Peizo1* mice reported by others (*69*).

Our studies using reporter mice that express endogenous PIEZO1 tagged with the red fluorescent protein tdT revealed that these channels transiently converge at sites where motile MΦs encounter one another. PIEZO1 channels were not observed to coalesce at sites where MΦs interacted with inert, stationary objects. Based on these findings, it is tempting to speculate that PIEZO1 channels may help to choreograph the regular distribution patterns observed in tissue resident MΦs in almost all tissues (*2*). Loss of PIEZO1, however, did not affect MΦ distribution in any of the tissues examined, suggesting that other factors are involved in homotypic cell sensing and the mutual repulsion of adjacent MΦs. Collectively, our data suggest that PIEZO1 has non-essential or redundant roles in MΦs under normal physiological conditions, distinct from its reported roles in non-physiological or diseased conditions (*34*, *61*, *62*).

## Acknowledgements

*Piezo1-tdTomato* mice were a kind gift from Prof Shaun Jackson (Heart Research Institute, University of Sydney). *Csf1r-Cre* mice were provided by Prof Edwin Hawkins (Walter and Eliza Hall Institute). *TdTomato-flox* mice were from Prof Ian Frazer (University of Queensland). We thank Dr Vaishnavi Ananthanarayanan (University of New South Wales) for kindly providing her expertise with imaging cells isolated from *Piezo1-tdTomato* mice on the Zeiss Elyra 7. We thank animal technicians at Australian BioResources, the University of New South Wales Animal Facility and the Laboratory Animal Core Facility at the Department of Biomedicine (Aarhus University). We acknowledge the expertise and input from staff at the Katharina Gaus Light Microscopy Facility (University of New South Wales) and the Bioimaging Core Facility at Aarhus University. Histology was provided by histologists at the Biotech Research Innovation Centre (BRIC, University of Copenhagen), Department of Dentistry (Aarhus University) and the Histology Core Facility at Biomedicine (Aarhus University). BioRender icons were utilized for figures in this manuscript. This work was funded by the National Health and Medical Research Council of Australia (1185021 to KP), the Novo Nordisk Foundation (NNF20OC0059705 to FMD) and the National Stem Cell Foundation of Australia (FMD). The μCT scanner was kindly donated by the VELUM foundation and some microscopes were kindly donated by the Carlsberg Foundation. DAH and KMI are grateful for funding support from the Mater Foundation.

---

**Fig S1:**
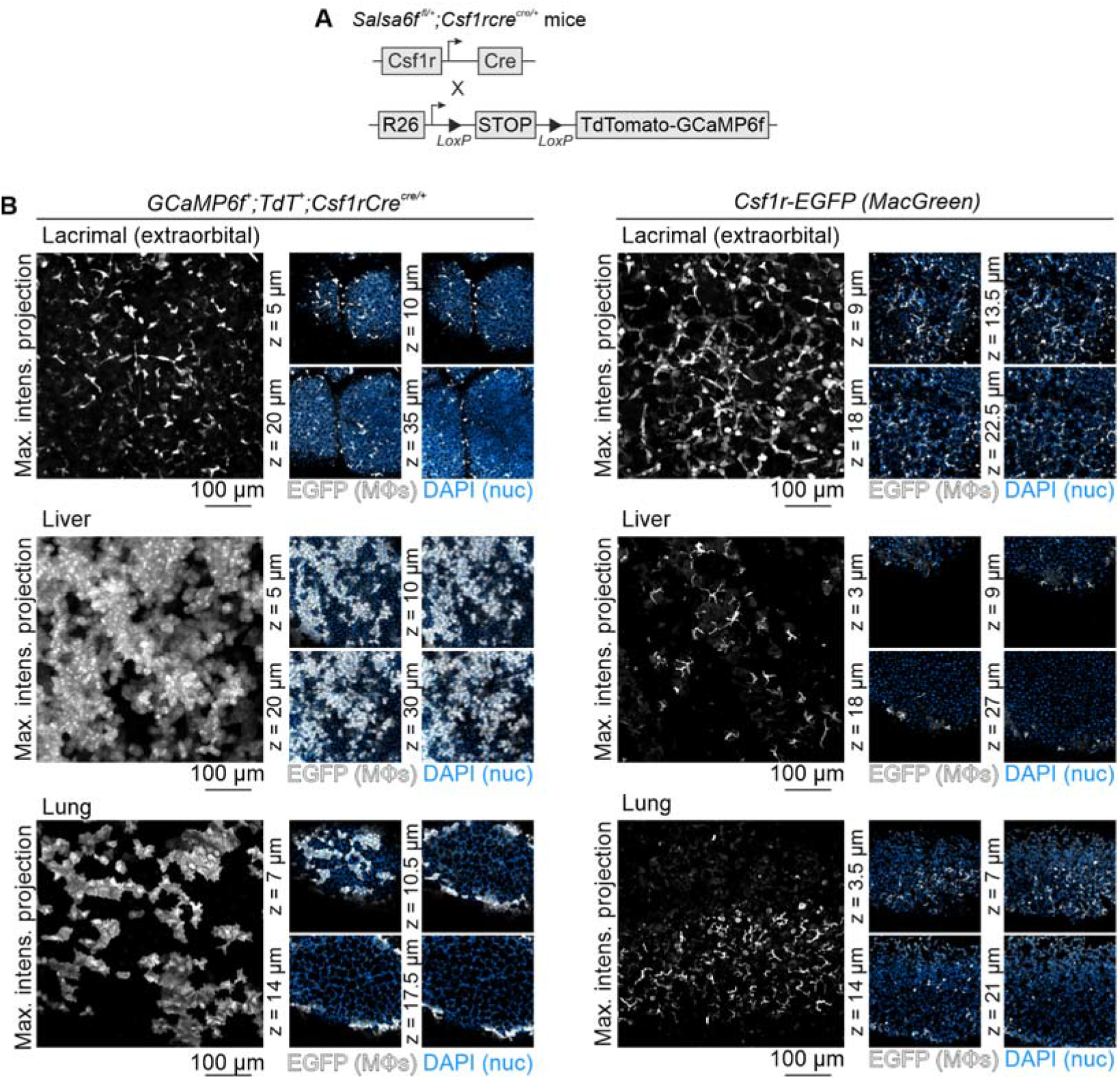
**A**, *GCaMP6f^+^;tdT^+^;Csf1r-cre^cre/+^*mice were generated by crossing *GCaMP6f-flox*, *tdTomato-flox* and *Csf1r-cre* mice (Fig 1) or by using *Salsa6f-flox* mice (containing tdT linked to GCaMP6f by a V5 epitope tag). **B**, Additional panel of organs imaged from *GCaMP6f^+^;tdT^+^;Csf1r-cre^cre/+^*vs *Macgreen* mice (related to Fig 1); *n* = 3.

**Fig S2:**
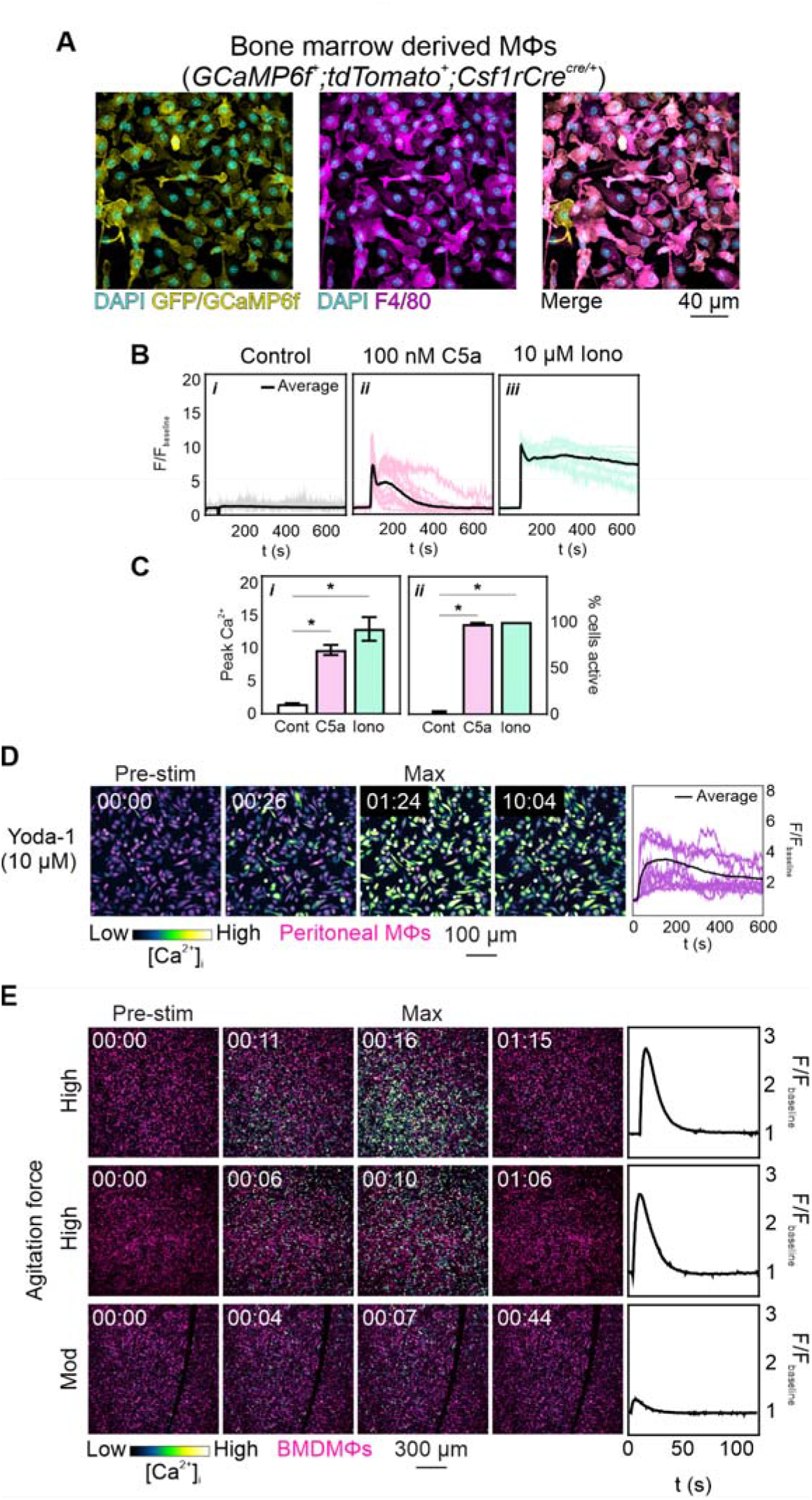
**A**, Bone marrow derived MΦs immunostained with anti-GFP/GCaMP6f (yellow) and anti-F4/80 (magenta) antibodies; nuclei are stained with DAPI (cyan). **B**, Intracellular Ca^2+^ responses to buffer only control, C5a (100 nM) and ionomycin (10 μM), quantified in **C**. Graphs show mean ± s.e.m., *n* = 3 independent experiments. Significance was assessed via ordinary one-way ANOVA with Tukey’s multiple comparison test, \**p* < 0.05 **D** Ca^2+^ responses in peritoneal MΦs treated with 10 µM Yoda1, representative of *n* = 2 independent experiments. **E** Intracellular Ca^2+^ responses of BMDMΦs monolayers flushed with inert buffer with high or moderate (mod) agitation force. Traces depict population Ca^2+^ responses. [min:sec].

**Fig S3:**
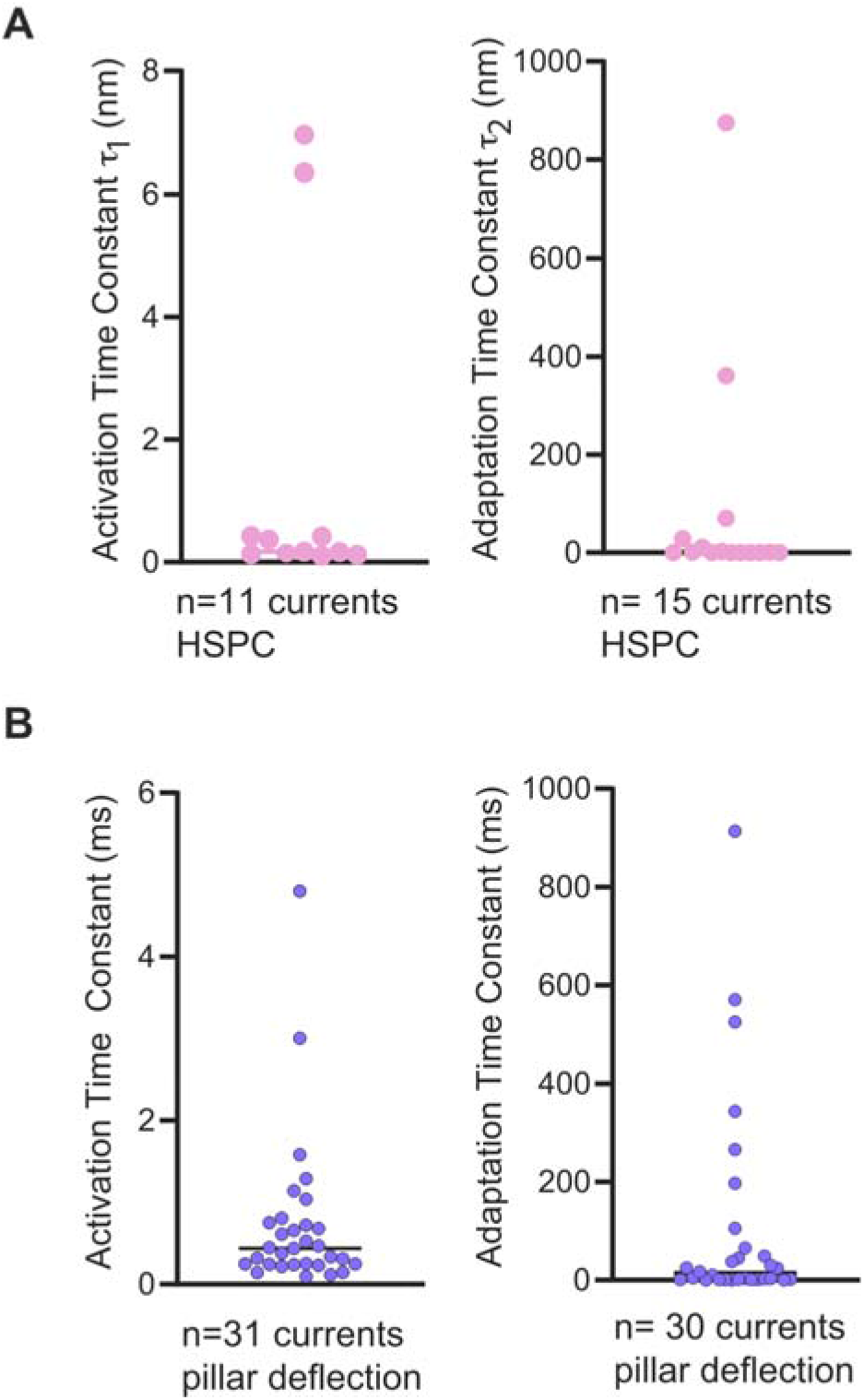
Activation and inactivation time constants for HSPC (**A**) and pillar deflection (**B**).

**Fig S4:**
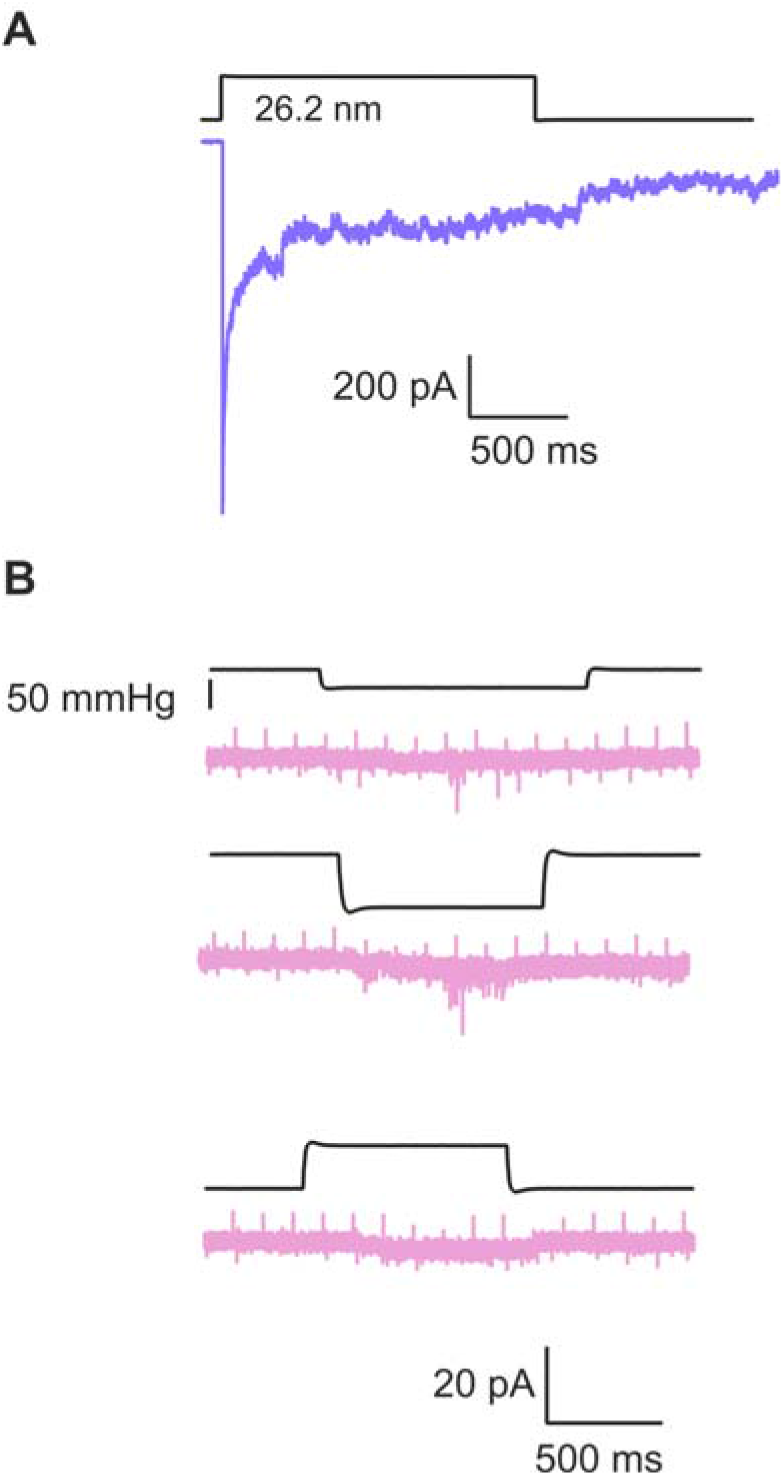
Pillar array and HSPC stimulation on a single cell. A MΦ, in whole-cell patch clamp mode, was initially stimulated by pillar deflection (**A**) which evoked a robust, sensitive current. An outside-out patch was then formed from the same cell and membrane stretch applied using HSPC (**B**).

**Fig S5:**
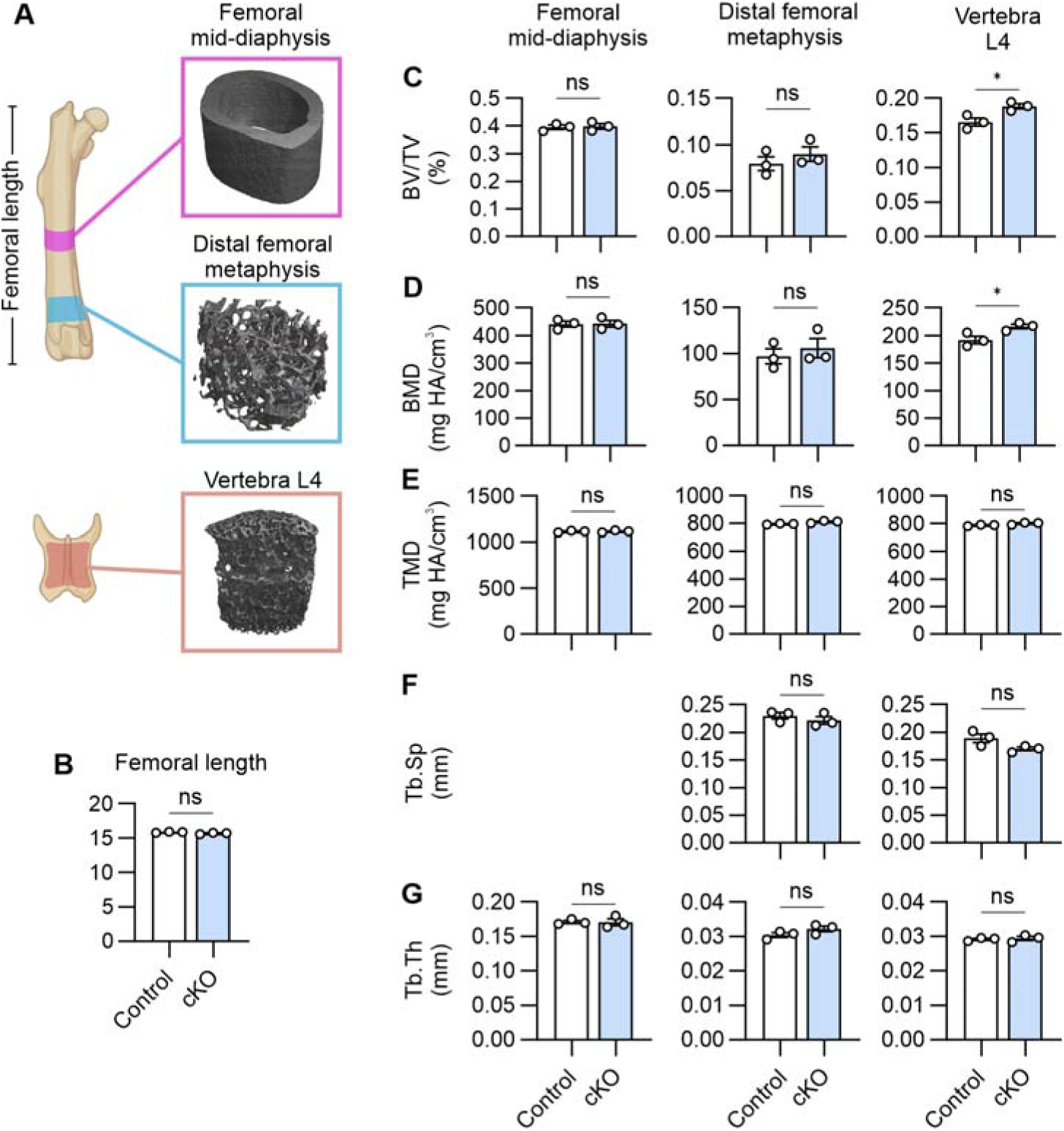
**A**, Schematic of bone regions scanned via microtomography, with example of scanned sample. Comparison of bone characteristics including femoral length (**B**), bone volume (BV)/total volume (TV) (**C**), bone mineral density (BMD), (**D**), tissue mineral density (TMD), (**E**), trabecular separation (Tb.Sp), (**F**), and trabecular thickness (Tb.Th), (**G**). Animals were post-lactational dams. Bars are mean ± s.e.m, *n* = 3 animals per cohort. Significance was assessed via unpaired t-test, \**p* < 0.05.

**Fig S6:**
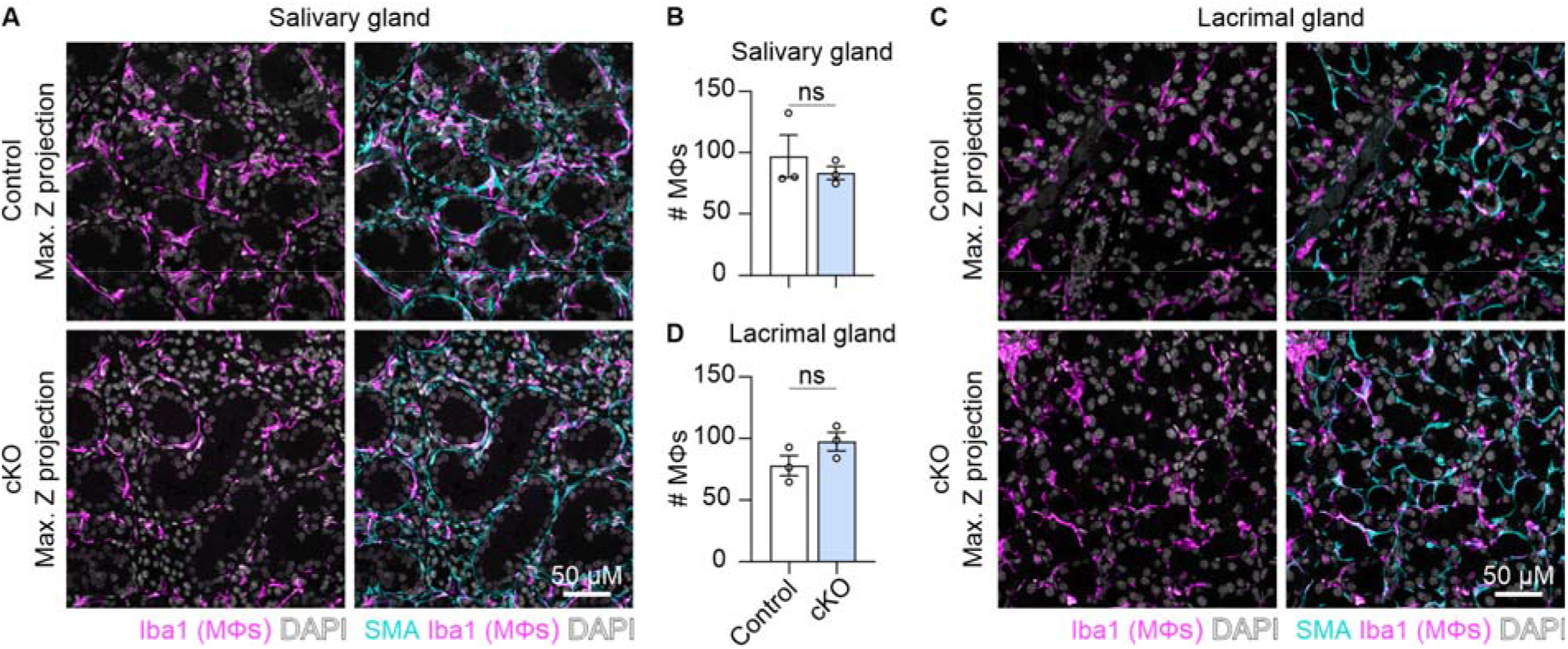
Immunohistochemical staining for smooth muscle actin (SMA, blue) and ionized Ca^2+^-binding adaptor molecule 1 (Iba1, magenta) in salivary gland (**A**) and lacrimal gland (**C**) isolated from cKO and control mice, with quantification of number of Iba1 positive cells in three randomly selected regions of each gland in **B** and **D**. Bars are ± s.e.m, individual data points are average value per animal. Data was analysed via unpaired t-test.

**Fig S7:**
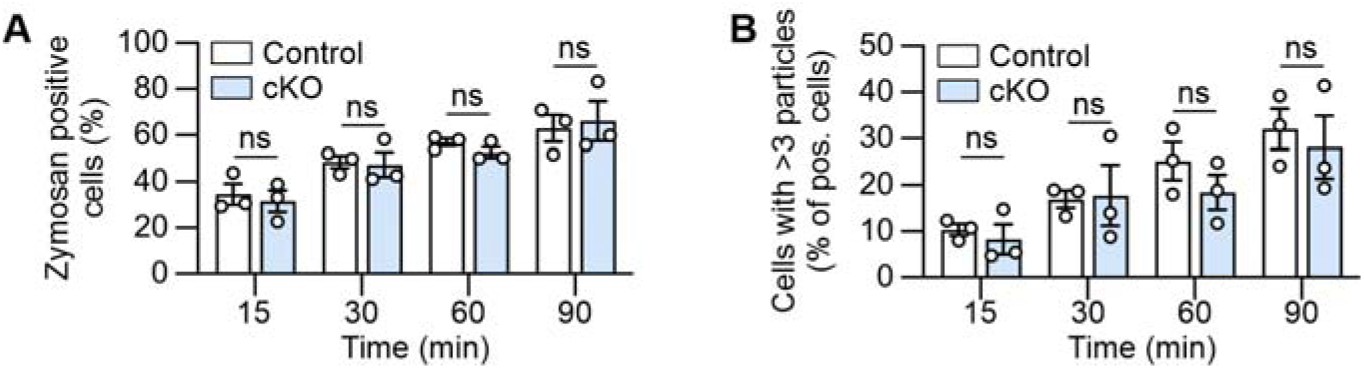
Comparison of zymosan uptake over time by peritoneal MΦs isolated from control and cKO animals. Bars are ± s.e.m, significance assessed via two-way ANOVA with Bonferroni post-test.

**Fig S8:**
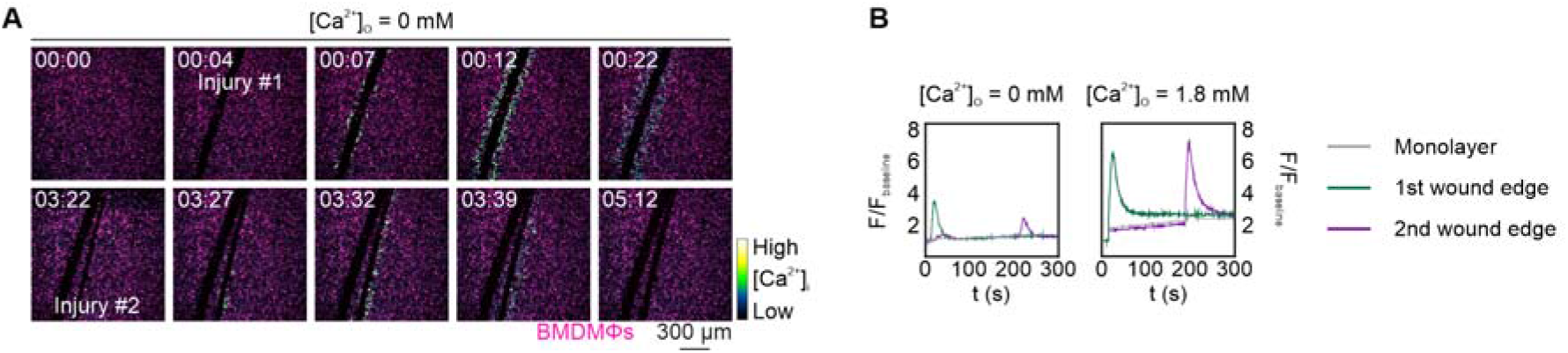
Ca^2+^ responses in BMDMΦ monolayers wounded under extracellular Ca^2+^ conditions. *n* = 3 independent experiments. [min:sec].

**Figure S9:**
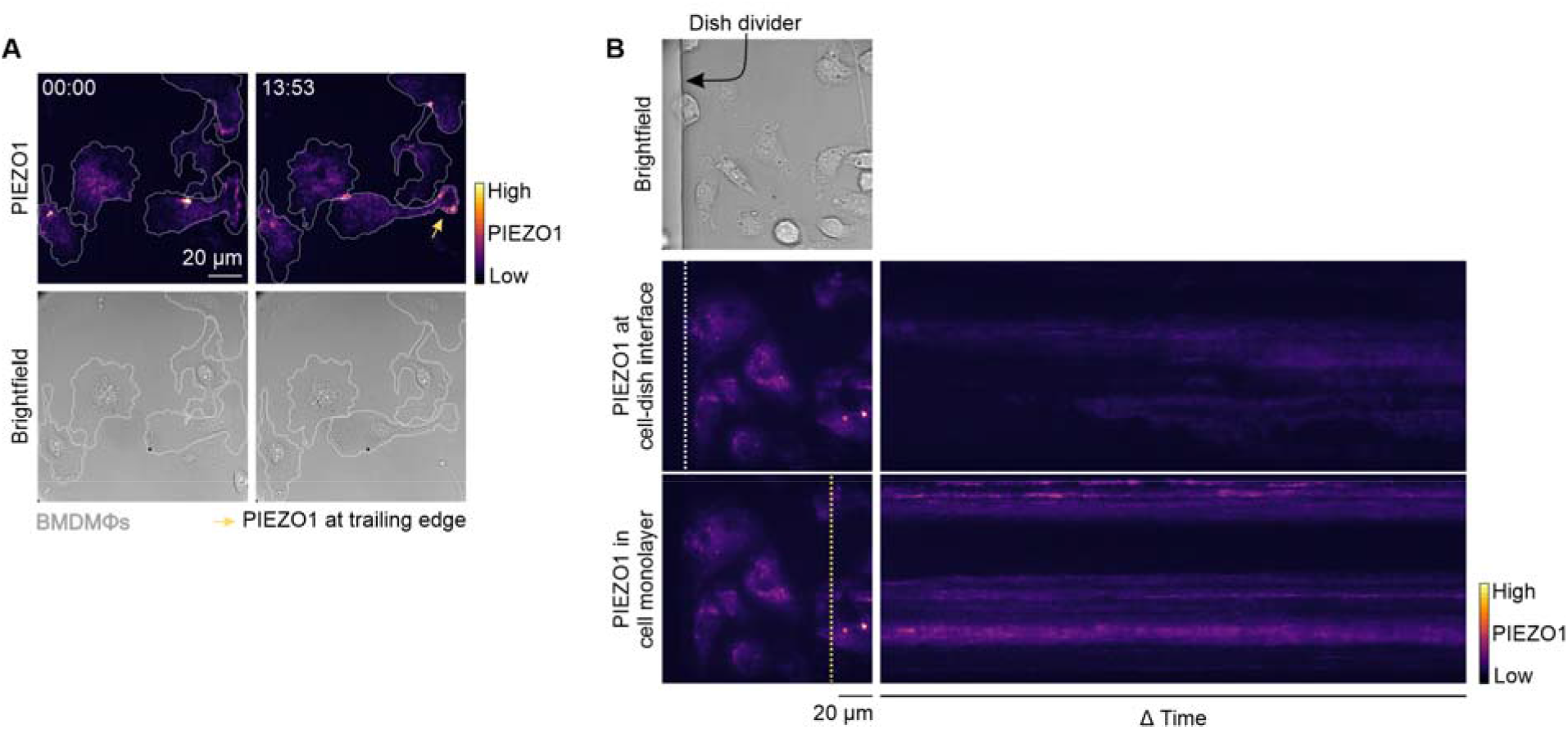
**A**, PIEZO1 channels coalesce at the trailing edge of some migrating cells, but rarely coalesce at contact sites with inert material (biocompatible silicone dish dividers), (**B**). [min:sec]

## Movie legends

**Movie S1**: Spontaneous Ca^2+^ activity in lymph node from *GCaMP6f^+^;tdTomato^+^;Csf1r-cre^cre^* mice. Recording shows 15 minutes, 232.96 µm × 232.96 µm. Representative of 3 independent experiments.

**Movie S2**: Mechanically invoked Ca^2+^ responses in ear skin from *GCaMP6f^+^;tdTomato^+^;Csf1r-cre^cre^*mice. Recording shows 14 min, 808.59 µm × 808.59 µm. Representative of 4 independent experiments.

**Movie S3**: Mechanically invoked Ca^2+^ responses in adipose tissue from *GCaMP6f^+^;tdTomato^+^;Csf1r-cre^cre^*mice. Recording shows 20 min 30 s, 931.82 µm × 931.82 µm. Representative of 4 independent experiments.

**Movie S4**: Ca^2+^-induced responses in peritoneal MΦs in response to zymosan particles. Recording shows 10 min 36 s, 212.13 µm × 212.13 µm. Representative of 4 independent experiments.

**Movie S5**: Ca^2+^ recording in response to physiological salt solution (PSS, control) in BMDMΦs. Text indicates time point of PSS addition, recording shows 9 min 30 s, 424.26 µm × 424.26 µm. Representative of 3 independent experiments.

**Movie S6**: Ca^2+^ responses to C5a (100 nM) in BMDMΦs. Text indicates time point of agonist addition, recording shows 10 min 3 s, 424.26 µm × 424.26 µm. Representative of 3 independent experiments.

**Movie S7**: Ca^2+^ responses to ionomycin (10 µM) in BMDMΦs. Text indicates time point of agonist addition, recording shows 8 min 3 s, 424.26 µm × 424.26 µm. Representative of 3 independent experiments.

**Movie S8**: Ca^2+^ responses to ATP (100 µM) in BMDMΦs, in the absence (left) or presence (right) of extracellular Ca^2+^ (1.8 mM). Text indicates time point of agonist addition, recording shows 10 min 3 s, 848.53 µm × 424.26 µm per side. Representative of 3 independent experiments.

**Movie S9**: Ca^2+^ responses to Yoda-1 (10 µM) in BMDMΦs. Text indicates time point of agonist addition, recording shows 14 min 15 s, 424.26 µm × 424.26 µm. Representative of 3 independent experiments.

**Movie S10**: Ca^2+^ responses to repeated pulses of fluid shear stress in BMDMΦs monolayer. Recording shows 4 min 50 s, 1272.79 µm × 1272.79 µm. Representative of 4 independent experiments; 1 with repeated flushing.

**Movie S11**: Ca^2+^ responses to mechanical wounding in BMDMΦs monolayer in the presence of extracellular Ca^2+^ (1.8 mM). Recording shows 5 min 19 s, 1272.79 µm × 1272.79 µm. Representative of 4 independent experiments.

**Movie S12**: Ca^2+^ responses to mechanical wounding in BMDMΦs monolayers with nominal extracellular Ca^2+^ present. Recording shows 5 min 19 s, 1272.79 µm × 1272.79 µm. Representative of 4 independent experiments.

**Movie S13**: BMDMΦ migratory behavior along a wound edge. Recording shows 17 h, 499.07 µm × 499.07 µm. Representative of 2 independent experiments.

**Movie S14**: PIEZO1 channels coalesce at sites of cell-cell contacts in patrolling BMDMΦs. Recording show 2 h 33 min 16 s, 78.39 µm × 78.39 µm. Representative of 4 independent experiments.

**Movie S15**: PIEZO1 channels coalesce at the trailing edge of BMDMΦs. Recording show 44 min 38 s, 123.88 µm × 123.88 µm. Representative of 4 independent experiments.

**Movie S16**: PIEZO1 channels in BMDMΦs rarely coalesce at inert dish divider edges. Recording show 46 min 12 s, 123.88 µm × 123.88 µm. Representative of 3 independent experiments.

## Methods

### Key Resource Table

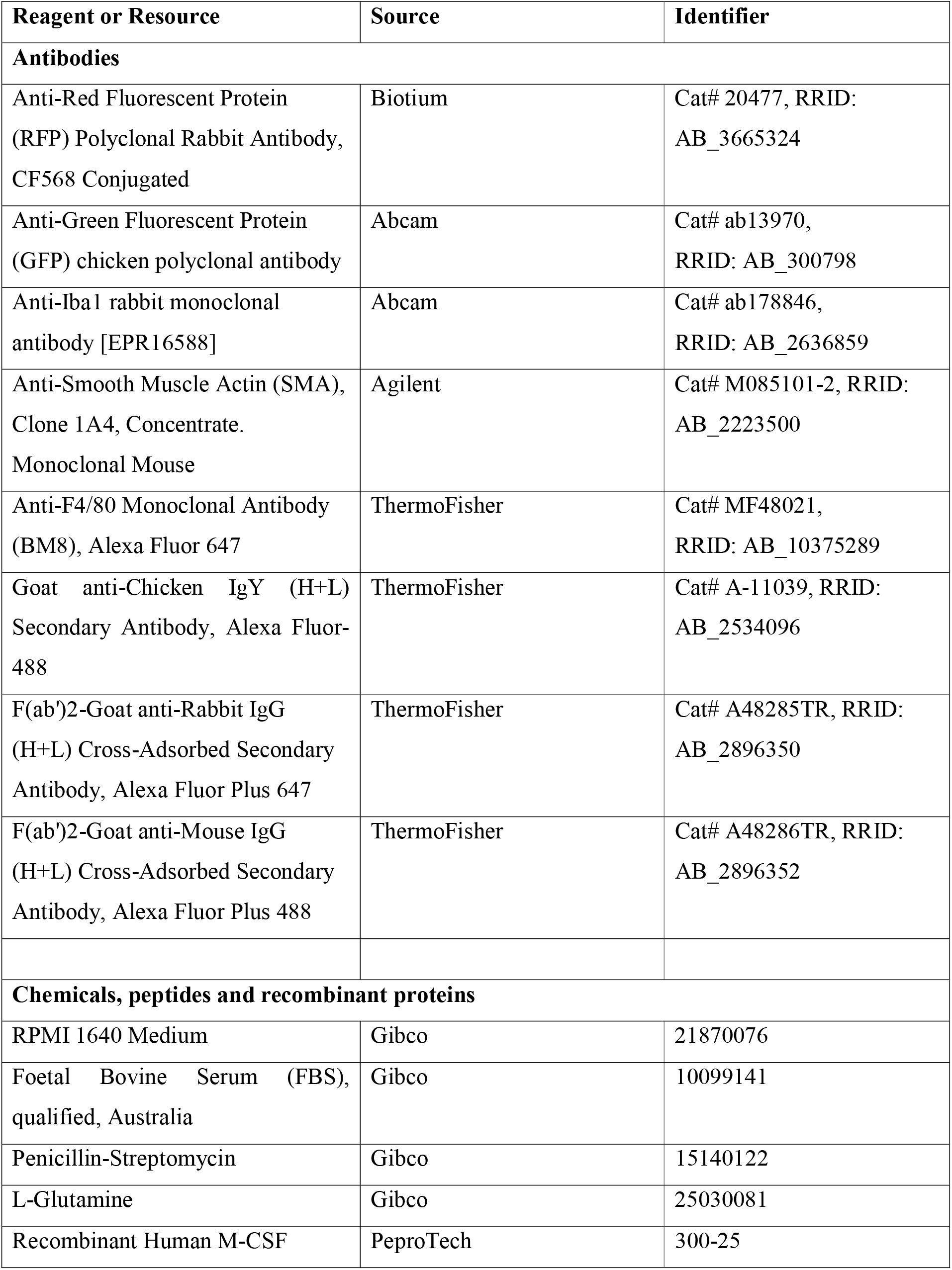
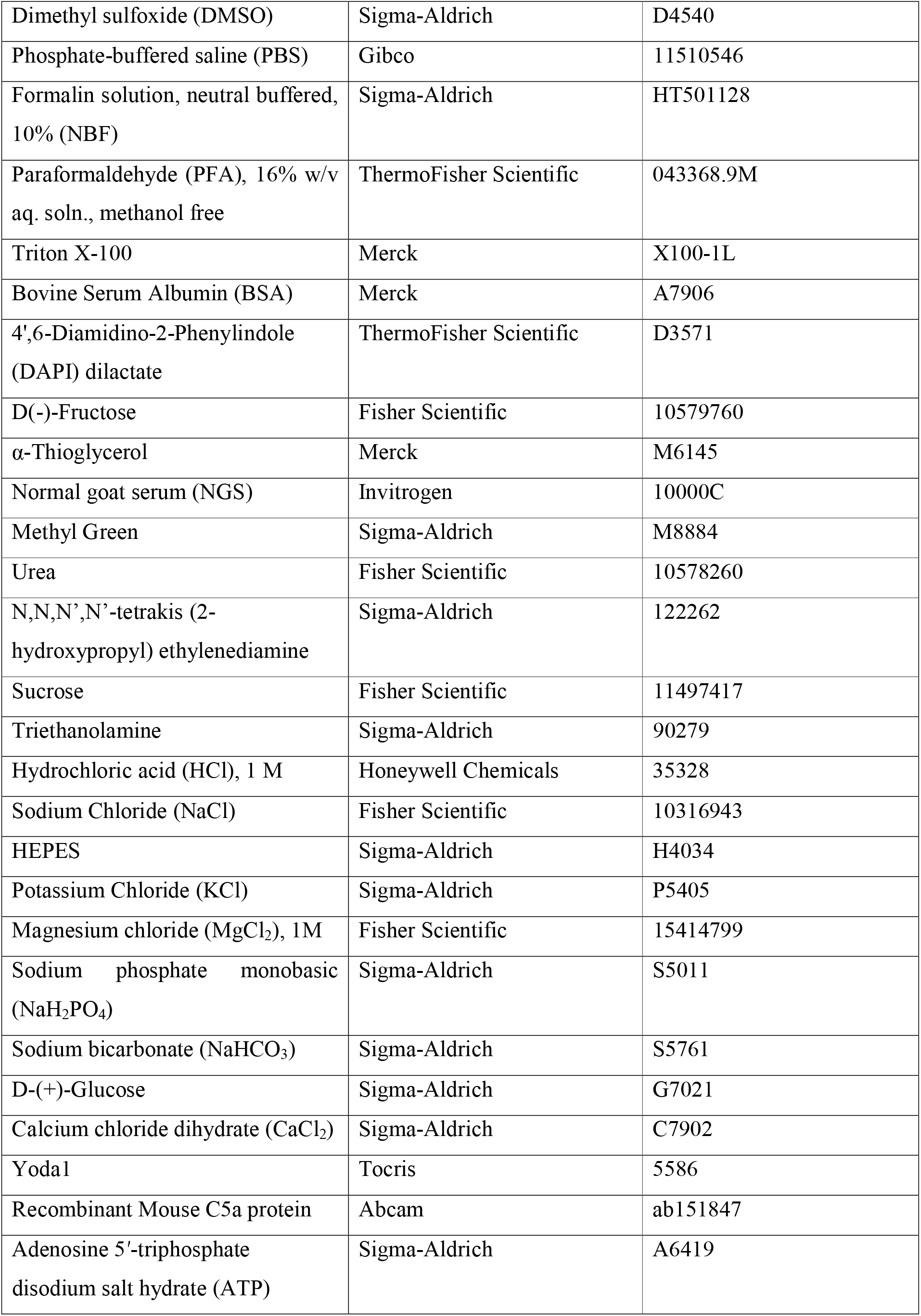
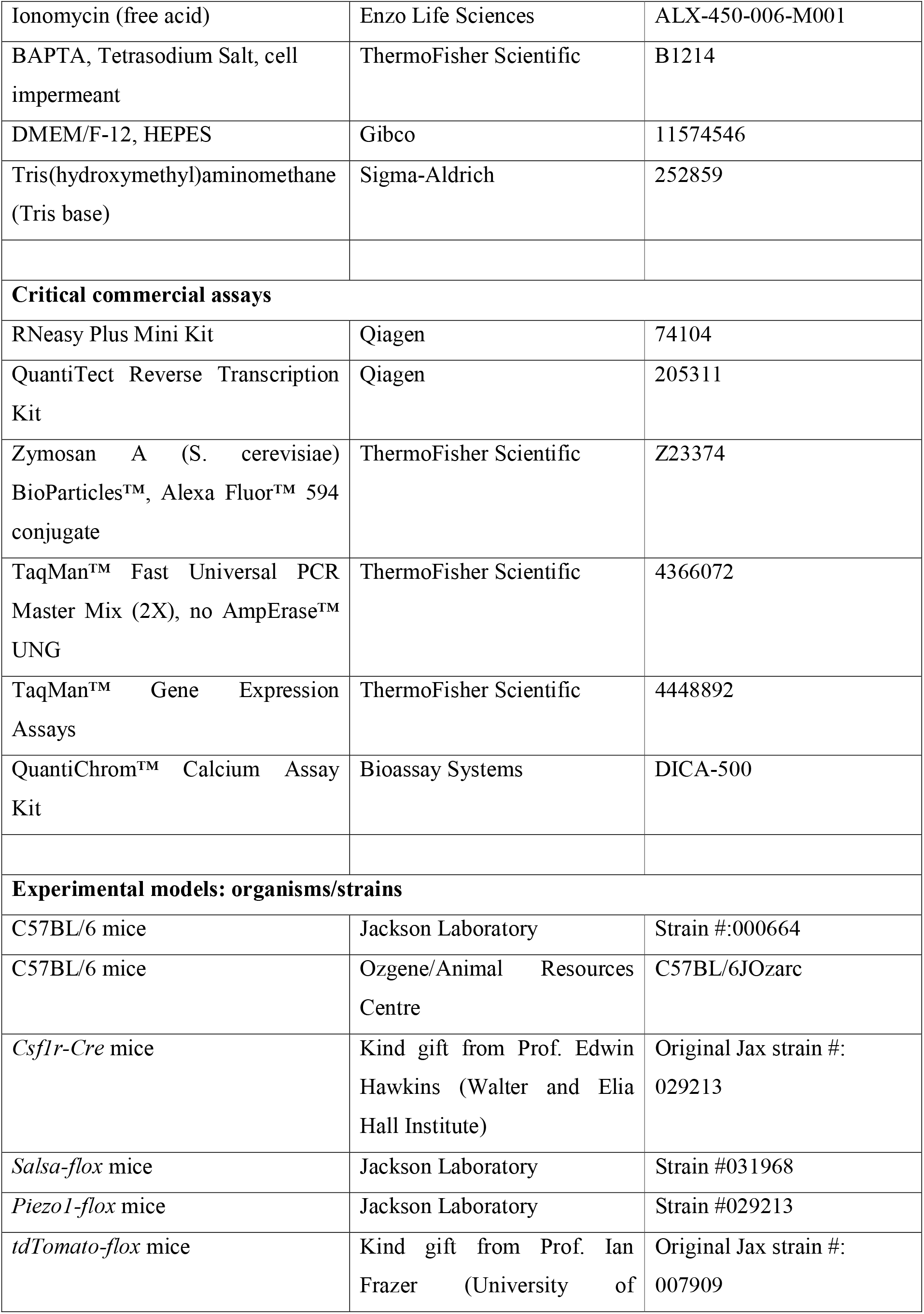
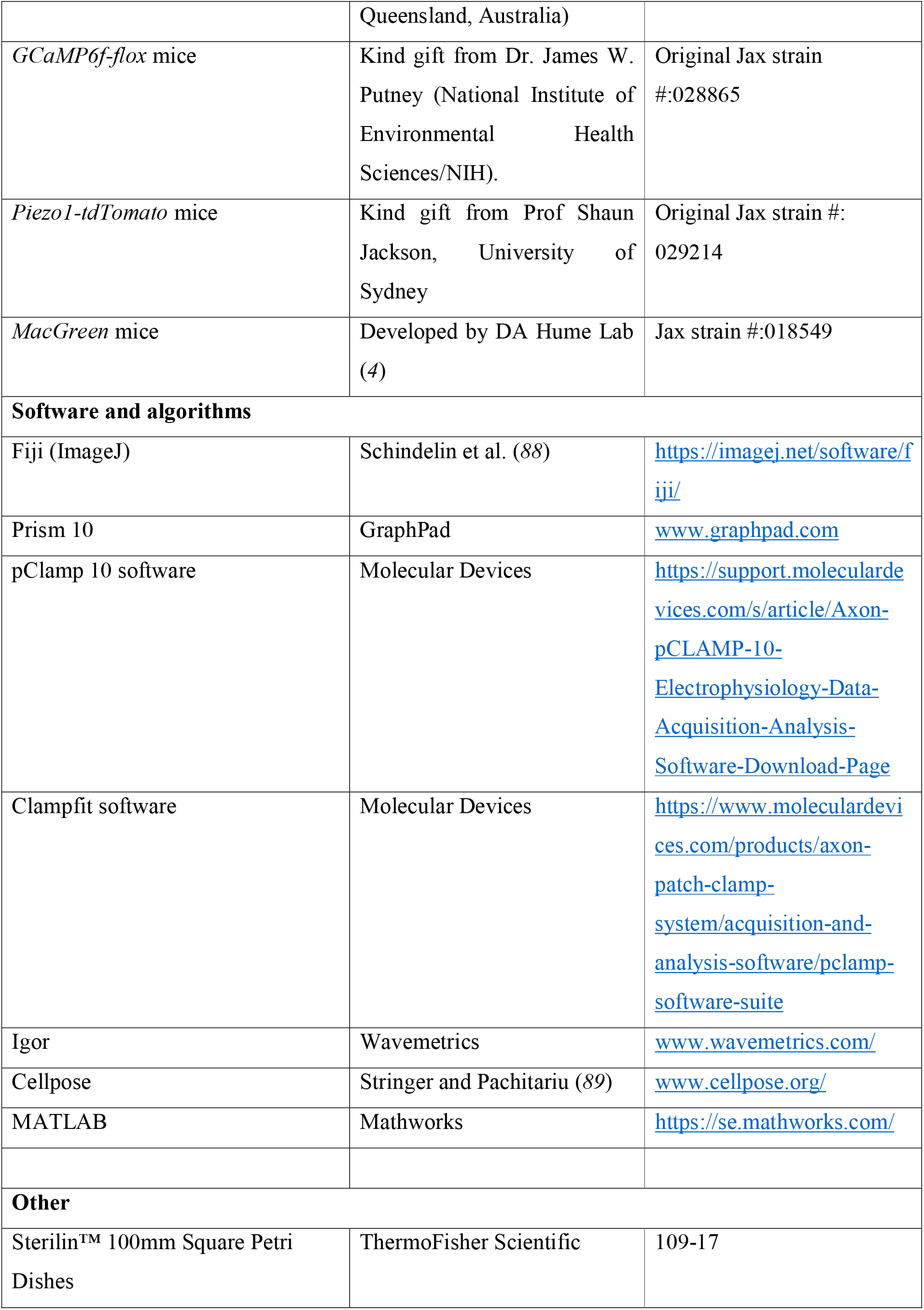
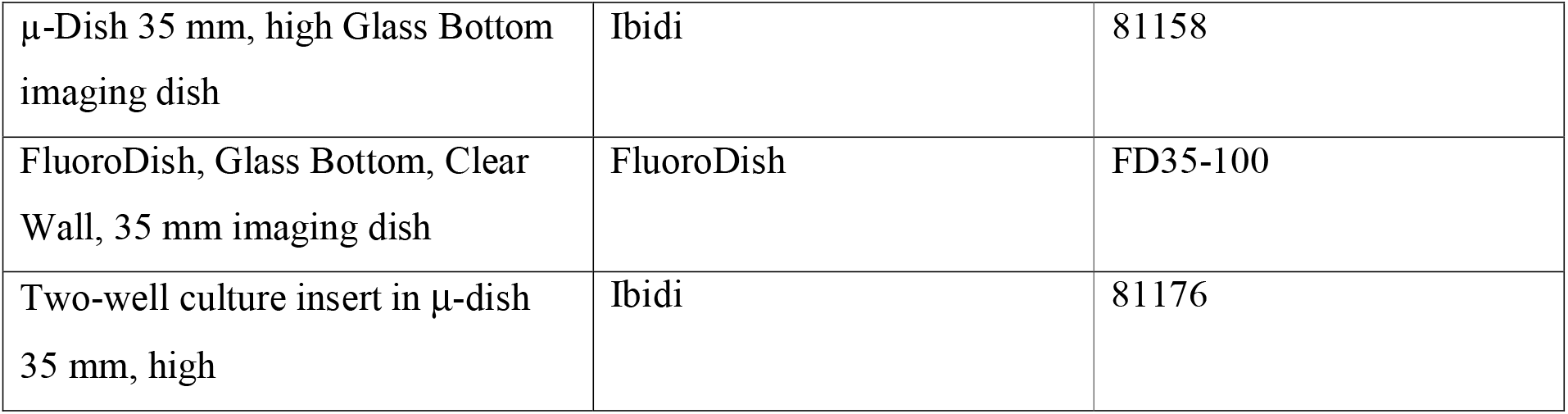

### Experimental model and subject details

#### Mice

Experiments were conducted in accordance with the Australian Code for the Care and Use of Animals for Scientific Purposes, the Queensland Animal Care and Protection Act (2001), the NSW Animal Research Act (1985), and the European Union Directive 86/609 with pre-approval from the Danish Animal Inspectorate (permit: 2022-15-0201-01196), with local animal ethics committee approvals. All strains were maintained on a C57BL6/J background. Animals were housed in specific pathogen free facilities, in individually ventilated cages with 12:12 h light-dark cycles, and *ad libitum* food and water. For histological comparisons, young mice were harvested between 10–14 weeks old, aged mice were ex-breeding females that were separated from breeding trios at 40 weeks and aged to 52 weeks prior to harvests. For lactation and involution studies, adult females were bred with C57BL6/J sires, allowed to litter naturally, and had litters standardised to 6–7 pups. Lactation tissue was collected during peak lactation (days 10–12 of lactation). For forced involution tissue, pups were weaned during peak lactation and tissue was harvested 168 h later (*90*).

### Method details

#### Cell culture

To generate BMDMΦs, mouse femurs were isolated and flushed twice with 5 mL cold PBS to extract bone marrow. Marrow was centrifuged for 5 min at 400 × *g* prior to resuspension in either freezing media (90% FBS and 10% DMSO) or MΦ cell culture media (10% heat-inactivated FBS, 2 mM L-glutamine, 100 U/mL Penicillin 100 µg/mL Streptomycin, 100 ng/mL Human M-CSF in RPMI 1640 medium). Cells were plated on Sterilin square petri dishes and maintained in a humidified cell culture incubator at 37°C, 5% CO_2_ for at least 7 days prior to use in experiments. Media was replenished after 3–4 days, if required. To replate BMDMΦs, monolayers were lifted from plasticware by flushing with an 18G needle. Peritoneal MΦs were isolated by injecting and removing 5 mL PBS in the peritoneal cavity. Recovered cell suspensions were centrifuged for 5 min at 400 × *g*, resuspended in MΦ cell culture media and plated onto imaging dishes. Dishes were incubated in a humidified cell culture incubator at 37°C, 5% CO_2_ for at least 2 h to allow attachment prior to use in experiments.

#### Tissue clearing

Tissues were fixed in 10% NBF for up to 6 hours, prior to storage in PBS at 4°C until sample processing. Tissues were cleared using the see deep brain (seeDB) protocol, as previously described (*91*, *92*). Briefly, tissue pieces were blocked and permeabilised in seeDB blocking buffer (1% w/v triton X-100 10% w/v BSA in PBS) at 4°C overnight. Samples were incubated in primary antibody dilutions at 4°C for 4 days, washed 3 x 1 h in PBS, then incubated in secondary antibody dilutions for a further 2 days prior to washing and nuclear DAPI staining. Following staining, samples were serially incubated in solutions of 20%, 40%, 60%, and 80% w/v fructose in distilled water (8–16 h incubation) prior to being submerged in 100% and 115% w/v fructose solutions (24 h incubation). All fructose solutions contained 0.5% v/v α-thioglycerol. Antibodies and DAPI dilutions were as follows: Anti-RFP-CF568 conjugated 1:300, Anti-GFP 1:1000, Anti-chicken-AF488 1:500, DAPI 10.9 µM. Cleared, immunostained tissue was imaged on an Olympus FV3000 laser scanning confocal microscope with UPLSAPO 10×/0.40, UPLSAPO 20×/0.75, UPLSAPO 30×/1.05 and UPLFLN 40×/0.75 objective lenses. Visualization and image processing was performed in ImageJ (v1.52e, National Institutes of Health) (*88*, *93*). Denoising of three-dimensional images was performed as described previously (*94*).

#### Immunofluorescence staining

Samples were stained as previously described (*56*), with minor adjustments. Briefly, cells were fixed for 15 min in 4% PFA prior to washing in PBS, and permeabilisation for 5 min in PBS with 0.1% Triton X-100. Samples were blocked for 1 h in blocking buffer (5% NGS, 0.05% Triton X-100 in PBS), prior to overnight incubation at 4°C with primary antibodies diluted in blocking buffer. Samples were then washed 3 × 10 min, then incubated with secondary antibodies diluted in blocking buffer for 1 h at room temperature. Following 3 × 10 min washes, samples were stained for 10 min with DAPI, washed a 2 × 10 min in PBS and then imaged. Antibodies and DAPI dilutions were as follows: Anti-RFP-CF568 conjugated 1:500, Anti-F4/80-AF647 conjugated 1:150, Anti-GFP chicken 1:1000, Anti-chicken-AF488 1:500, DAPI 2.18 µM. Samples were imaged on an Olympus FV3000 laser scanning confocal microscope as described above, or via a Leica SP8 confocal microscope with a 40× 1.3 HC PL APO CS2 Oil objective lens. Image visualization and processing was conducted as described above.

#### Histology and immunohistochemistry

Tissues were fixed, processed, and paraffin embedded as previously described (*90*, *95*). Formalin-fixed paraffin-embedded slides were sectioned to 4–5 µm, deparaffinized by immersion in xylene (3 × 5 min immersions) and rehydrated via a reducing ethanol series. Sections were permeabilised for 5 min in 0.5% Triton X-100 in PBS. Antigen retrieval was performed in a Decloaking Chamber NxGen digital pressure system (Biocare Medical) at 110°C for 11 min, in a sodium citrate buffer (0.01 M, pH 6). Samples were blocked for 1 h in a humidified chamber at room temperature in blocking buffer (10% NGS, 0.05% Triton X-100 in PBS). Primary antibodies were diluted in antibody dilution buffer (5% NGS, 0.025% Triton X-100 in PBS) and incubated on sections at 4°C in a humidified chamber overnight. Following washing (3 × 5 min), sections were incubated in secondary antibodies diluted in antibody dilution buffer for 1 h in a humidified chamber at room temperature. Sections were washed 3 × 5 min, stained with DAPI for 10 min, washed a further 3 × 5 min, mounted in 50:50 glycerol:PBS, and imaged within 48 h. Antibodies and DAPI dilutions were as follows: Anti-IBA1 rabbit 1:600, Anti-SMA mouse 1:600, Anti-rabbit-AF647 1:500, Anti-mouse AF488 1:500, DAPI 1.36 µM. Samples were imaged via a Leica STELLARIS 8 with a HC PL APO 40×/1.25 glycerol motorised correction collar CS2 objective. Three regions were imaged per tissue per animal; regions were chosen via DAPI channel only. Number of MΦs per tissue were manually counted by a blinded analyst.

#### Wholemount histology

Inguinal mammary glands were dissected, spread on Tetra-Pak card and fixed in 10% NBF overnight in room temperature as previously described (*90*, *91*). Post fixing, samples were immersed in CUBIC reagent 1 (25% w/w urea, 25% w/w N,N,N’,N’-tetrakis (2-hydroxypropyl) ethylenediamine, 15% w/w Triton X-100 in distilled H2O) for 3 days at 37°C, in a humidified chamber, with once daily reagent refreshment and sample inversion (*91*, *96*). Glands were stained in 0.5% methyl green solution for 1.5 h at room temperature, with gentle rocking, washed 3 times in water and then destained for 20 min in 50% ethanol with 25 mM HCl. Samples were incubated in modified CUBIC reagent 2 (sucrose 50% w/w, urea 25% w/w, triethanolamine 10% w/v, Triton X-100 0.1% w/v, 25 mM NaCl in distilled water) at 37°C in a humidified chamber overnight before imaging. Samples were imaged on a Leica M205FCA stereo microscope with a planapo 1.0× M series objective.

#### Real-time reverse transcription PCR

RNA was isolated from BMDMΦs via Qiagen Buffer RLT Plus, prior to purification with a RNeasy Plus Mini Kit. Reverse transcription was conducted with using the Quantitect Reverse Transcription Kit and resulting cDNA was amplified on a StepOne Plus Real Time PCR system, using Taqman Fast Universal PCR Master Mix and Taqman Gene Expression assays. Relative quantitation was calculated with reference to *Hprt* RNA and analysed using the comparative C_T_ method (*97*). C_T_ values above 35 were set to 35 were considered undetected. values The following assays were used in this study: *Hprt* Mm03024075_m1*, Trpv4* Mm00499025_m1*, Piezo1* Mm01241549_m1*, Piezo2* Mm01265861_m1*, Trpv2* Mm00449223_m1*, Trpm7* Mm00457998_m1*, P2rx4* Mm00501787_m1*, P2rx7* Mm01199500_m1*, Orai1* Mm00774349_m1*, Stim1* Mm01158413_m1.

#### Live cell Ca^2+^ imaging

All live cell Ca^2+^ imaging was conducted at room temperature in physiological salt solution (PSS: 10 mM HEPES, 5.9 mM KCl, 1.4 mM MgCl_2_, 1.2 mM NaH_2_PO_4_, 5 mM NaHCO_3_, 140 mM NaCl, 11.5 mM glucose and 1.8 mM CaCl_2_; pH 7.3–7.4). Extracellular Ca^2+^ free experiments were conducted in PSS without the addition of 1.8 mM CaCl_2_ and the presence of 500 µM BAPTA. For studies assessing responses to pharmacological stimuli, cells were bathed in PSS and received either buffer control (PSS), or agonists at concentrations indicated in figure legends. Zymosan uptake experiments were conducted using particles at a concentration of 0.2 mg/mL. Live cell imaging was conducted on either an Olympus FV3000 laser scanning confocal microscope with UPLSAPO 10×/0.40, UPLSAPO or UPLSAPO 30×/1.05 objective lenses, or on a Leica STELLARIS 8 with a HC FLUOTAR L 25×/0.95 W VISIR objective.

#### Ca^2+^ responses in live tissue

Ear and adipose tissues were harvested, cut into 3–5 mm^2^ pieces and incubated in DMEM/F12 containing 10% FBS in a 37°C humidified cell culture incubator with 5% CO_2_ until ready for imaging as previously described (*98*, *99*). Inguinal lymph nodes were removed from surrounding tissue and stored in DMEM/F12 at 37°C until ready for imaging. Immediately prior to imaging, tissue pieces were placed on an imaging dish and bathed in PSS. Mechanically induced Ca^2+^ recordings in ear and adipose tissue were created by manually prodding tissue pieces with an 18G microlance needle. Imaging was conducted either on a Leica M205FCA stereo microscope with a planapo 1.0× M series objective (Figure 1E) or on a Leica STELLARIS 8 with a HC PL APO 10×/0.40 CS2 objective (Figure 1D&F).

#### Live cell imaging of wounded BMDMΦ monolayers

Wild-type BMDMΦ monolayers were prepared as described above, plated in a 96-well plate and wounded via manual scratching via a 200 µL pipette tip. Wounded monolayers were imaged at 37°C overnight in a PhaseFocus Livecyte live cell imaging system.

#### Flow cytometry

GCaMPP6f signal in BMDMΦs was assessed during flow cytometry using a Beckman Coulter CytoFLEX-S. Briefly, cells were suspended in either ethanol or a flow cytometry buffer (PBS/2% FBS). During sorting, BMDMΦs in buffer were treated with 10 µM Yoda1 after 200 sec, and the flow rate was subsequently increased from 10 µL/minute to 60 µL/minute after 300 sec.

#### Piezo1-tdTomato live cell imaging

Prior to imaging, *Piezo1^tdT^* BMDMΦs were isolated, differentiated, and replated as described above. Following replating, monolayers were maintained in MΦ media without penicillin/streptomycin. Prior to imaging, cell monolayers were scratched with a 10 µL pipette tip. Cells were imaged at 37°C at a TIRF angle of 63.5, using a Zeiss ELYRA 7 microscope with a 63× Plan Apochromat oil immersion objective with 1.46 N.A.

#### Image analysis

Supplementary movies were prepared in FIJI. Calcium channel LUT was set as either mpl-inferno (Movies S1-3) or Green Fire Blue (Movies S4-12).

For monolayer Ca^2+^ agonist response data, the first 10 images of the tdT channel were summed in FIJI and resulting image had individual cells segmented via Cellpose (v3.1.1.2). Resulting masks were transferred to GCaMP6f channel in FIJI for parameter measurement over time. Recorded values were analysed in MATLAB (R2021b). Fluorescence values over time were normalised to the mean fluorescence value for the first 10 images for each individual cell (F/F_baseline_). Average peak Ca^2+^ levels were determined by averaging maximum F/F_baseline_ values for all individual cell responses per dish, statistics were performed on dish averages. Cell activation was defined as the F/F_baseline_ value of a cell exceeding a threshold fluorescence level during imaging time (≥2 F/F_baseline_ for responses recorded on Olympus FV3000 and ≥3 F/F_baseline_ for responses recorded on Leica STELLARIS8). Individual traces for plots were randomly selected. For fluid flow monolayer Ca^2+^ response data, individual cell segmentation was not conducted due to frequent dish movements during imaging periods, instead whole dish fluorescence was recorded and analysed using the methods described above. For monolayer scratch analysis, 10 images post wounding were summed in FIJI in the tdT channel were and the resulting image had individual cells segmented via Cellpose. Resulting masks were transferred to GCaMP6F channel in FIJI and measured as described above. Scratch wounds were segmented by hand in Cellpose, and resulting masks was transferred to FIJI to permit measurement of perimeter points. Cell fluorescence values were analyzed in MATLAB as described above. The Euclidean distance between cell centroids and each perimeter point of scratch wounds was calculated to determine the shortest distance between each cell and wound edges. Cells within 90 µm of wound edges were classified as ‘wound edge’ cells, cells further than 90 µm from any wound edge were classified as ‘monolayer’ cells.

#### Combined mechanical stimulation and electrophysiological recordings

Two distinct methods were utilised to apply mechanical stimuli to delimited regions of the membrane, a pillar array assay to apply mechanical inputs directly at cell-substrate contact points and high speed pressure clamp (HSPC) to stretch a region of membrane within an outside-out patch. Pipettes for whole-cell patch clamp and outside-out patch clamp analyses were prepared using a pipette puller fitted with a box filament (P-1000, Sutter Instruments, USA) from thick-walled filamented glass (Harvard Apparatus, USA). Before use, a homemade micro-forge was utilised to heat-polish the glass, resulting in a final resistance of 3–6 MΩ. For measurements, the internal solution contained 110 mM KCl, 10 mM NaCl, 1 mM MgCl2, 1 mM EGTA and 10 mM HEPES (pH 7.3) and the extracellular solution 140 mM NaCl, 4 mM KCl, 2 mM CaCl2, 1 mM MgCl2, 4 mM glucose and 10 mM HEPES (pH 7.4). Data were obtained using a Nikon Ti-E inverted microscope and an Axopatch 200B with pClamp 10 software (Molecular Devices, USA). Pipette and membrane capacitance were compensated and to minimise voltage errors, series resistance was compensated by at least 60%. Mechanically-activated currents were recorded at a holding potential of −60 mV. Patch clamp data were subsequently analysed using Clampfit software (Molecular Devices, USA).

Two distinct methods were utilised to apply mechanical stimuli to delimited regions of the membrane. A pillar array assay was used to apply mechanical stimuli at connections between the MΦs and the substrate, as previously described (*73*, *100*). Cells were cultured on uncoated pillar arrays, activated using oxygen plasma (Diener GmbH, Germany), before exchange of media for extracellular buffer. After establishing whole-cell patch clamp configuration, mechanically activated currents were recorded in voltage clamp mode while stimuli were applied by serially deflecting an individual pilus subjacent to the cell. The pillar deflection was achieved using a blunt, heat-polished pipette (tip diameter approx. 2 µm) driven by a MM3A-LS nanomanipulator (Kleindiek Nanotechnik, Germany) and positioned adjacent a pilus. Multiple stimuli ranging between 1 nm–1 µm were applied with a delay between each stimulus of at least 10 s. Give the propensity of MΦs to engulf the pillar structures subjacent to the cell, the tdT channel was used to select stimulation sites where the cell membrane was covering but not engulfing the subjacent pilus. In addition, small stimuli adjacent, but not in contact with, the pilus were applied and data were discarded if currents were mechanically evoked in the absence of pillar movement. Data were also discarded if the current trace did not return to baseline after the end of the pillar deflection. To quantify the stimulus, bright-field images of the deflected pilus were taken before and during each stimulation using a 40× objective (NA 0.6). The centre of each pilus was calculated in a post-hoc analysis by applying a 2D-gaussian fit of intensity values in the bright field visualisation of each pilus (Igor, Wavemetrics, USA) and the magnitude of pillar deflection calculated from successive images by comparing the difference in the calculated centre point.

Membrane stretch stimuli were applied to outside-out patches of membrane using high speed pressure clamp (HSPC). After formation of a high resistance seal and establishing whole-cell mode, an outside-out patch was formed from extraction of a region of membrane. Both positive and negative pressure stimuli were then applied, with at least 10 s intervals between each stimulus, while currents traces were recorded in voltage clamp mode, at a holding potential of −60 mV.

#### Milk collection and analysis

Milk samples were collected from lactating dams as previously described (*90*). Briefly, pups were removed from dams for up to 2 hours prior to milking. Dams were administered 2 IU of oxytocin via intraperitoneal injection, then lightly anesthetized via 2% isoflurane. Milking was conducted on inguinal mammary glands of both the left and right side through manual manipulation of the glands. Expressed milk was collected from the nipple tip by pipette. Collected samples were stored at −20°C until analysis. Free milk Ca^2+^ concentrations were measured using a phenolsulphonephthalein reaction with a QuantiChrom Calcium Assay Kit as per manufacturer’s instructions.

#### Bone sample analysis

Femura and L4 vertebrae were harvested and stored at −20°C in Ringer’s solution (2.5 mM Tris base, 154 mM NaCl, 1.63 mM CaCl_2_, 5.63 mM KCl, pH 7.4) until analysis. The microarchitectural bone parameters were determined using a desktop µCT scanner (Scanco µCT 35, Scanco Medical AG, Brüttiselen, Switzerland) as previously described (*101*). In brief: Scans were performed with 1000 projections/180°, isotropic resolution of 3.5 µm, X-ray tube voltage of 55 kV and current of 114 µA, and an integration time of 800 ms. Subsequently, the microstructural parameters were determined using IPL (v 5.11, Scanco Medical AG, Brüttiselen, Switzerland). Femoral bone length was determined using a digital sliding caliper, measured from the top of the femoral head.

#### Statistical analysis

Statistical analysis was performed using GraphPad Prism (v10.1.0). Statistical tests are outlined in figure legends. All analyses were conducted on average values for independent repeats, where *n* = 3 to 6 animals, as indicated.

## References

1. D. Hume, K. Irvine, C. Pridans, The Mononuclear Phagocyte System: The Relationship between Monocytes and Macrophages. Trends in Immunology 40, 98–112 (2019).

2. E. Mass, F. Nimmerjahn, K. Kierdorf, A. Schlitzer, Tissue-specific macrophages: how they develop and choreograph tissue biology. Nat Rev Immunol 23, 563–579 (2023).

3. S. J. Jenkins, D. A. Hume, Homeostasis in the mononuclear phagocyte system. Trends Immunol 35, 358–367 (2014).

4. R. T. Sasmono, D. Oceandy, J. W. Pollard, W. Tong, P. Pavli, B. J. Wainwright, M. C. Ostrowski, S. R. Himes, D. A. Hume, A macrophage colony-stimulating factor receptor-green fluorescent protein transgene is expressed throughout the mononuclear phagocyte system of the mouse. Blood 101, 1155–63 (2003).

5. D. A. Ovchinnikov, W. J. M. van Zuylen, C. E. E. DeBats, K. A. Alexander, S. Kellie, D. A. Hume, Expression of Gal4-dependent transgenes in cells of the mononuclear phagocyte system labeled with enhanced cyan fluorescent protein using Csf1r-Gal4VP16/UAS-ECFP double-transgenic mice. J Leukoc Biol 83, 430–433 (2008).

6. T. A. Stewart, K. Hughes, D. A. Hume, F. M. Davis, Developmental Stage-Specific Distribution of Macrophages in Mouse Mammary Gland. Frontiers in Cell and Developmental Biology 7 (2019).

7. F. Rae, K. Woods, T. Sasmono, N. Campanale, D. Taylor, D. A. Ovchinnikov, S. M. Grimmond, D. A. Hume, S. D. Ricardo, M. H. Little, Characterisation and trophic functions of murine embryonic macrophages based upon the use of a Csf1r-EGFP transgene reporter. Dev Biol 308, 232–246 (2007).

8. E. S. Nakasone, H. A. Askautrud, T. Kees, J. H. Park, V. Plaks, A. J. Ewald, M. Fein, M. G. Rasch, Y. X. Tan, J. Qiu, J. Park, P. Sinha, M. J. Bissell, E. Frengen, Z. Werb, M. Egeblad, Imaging tumor-stroma interactions during chemotherapy reveals contributions of the microenvironment to resistance. Cancer Cell 21, 488–503 (2012).

9. A. Sehgal, D. S. Donaldson, C. Pridans, K. A. Sauter, D. A. Hume, N. A. Mabbott, The role of CSF1R-dependent macrophages in control of the intestinal stem-cell niche. Nature Communications, doi: 10.1038/s41467-018-03638-6 (2018).

10. E. Mass, I. Ballesteros, M. Farlik, F. Halbritter, P. Günther, L. Crozet, C. E. Jacome-Galarza, K. Händler, J. Klughammer, Y. Kobayashi, E. Gomez-Perdiguero, J. L. Schultze, M. Beyer, C. Bock, F. Geissmann, Specification of tissue-resident macrophages during organogenesis. Science 353, aaf4238 (2016).

11. S. J. Jenkins, J. E. Allen, The expanding world of tissue-resident macrophages. Eur J Immunol 51, 1882–1896 (2021).

12. M. J. Berridge, P. Lipp, M. D. Bootman, The versatility and universality of calcium signalling. Nature reviews. Molecular cell biology 1, 11–21 (2000).

13. R. Y. Tsien, New calcium indicators and buffers with high selectivity against magnesium and protons: design, synthesis, and properties of prototype structures. Biochemistry 19, 2396–2404 (1980).

14. R. Y. Tsien, T. J. Rink, M. Poenie, Measurement of cytosolic free Ca2+ in individual small cells using fluorescence microscopy with dual excitation wavelengths. Cell Calcium 6, 145–157 (1985).

15. G. Grynkiewicz, M. Poenie, R. Y. Tsien, A new generation of Ca2+ indicators with greatly improved fluorescence properties. J Biol Chem 260, 3440–3450 (1985).

16. T. H. Steinberg, A. S. Newman, J. A. Swanson, S. C. Silverstein, Macrophages possess probenecid-inhibitable organic anion transporters that remove fluorescent dyes from the cytoplasmic matrix. J Cell Biol 105, 2695–2702 (1987).

17. S. M. McMahon, M. B. Jackson, An Inconvenient Truth: Calcium Sensors Are Calcium Buffers. Trends Neurosci 41, 880–884 (2018).

18. M. D. Bootman, S. Allman, K. Rietdorf, G. Bultynck, Deleterious effects of calcium indicators within cells; an inconvenient truth. Cell Calcium 73, 82–87 (2018).

19. X. Li, Y.-X. Du, C.-L. Yu, N. Niu, Ion channels in macrophages: Implications for disease progression. Int Immunopharmacol 144, 113628 (2025).

20. T. A. Stewart, F. M. Davis, An element for development: Calcium signaling in mammalian reproduction and development. Biochimica et Biophysica Acta - Molecular Cell Research 1866 (2019).

21. M. Vaeth, I. Zee, A. R. Concepcion, M. Maus, P. Shaw, C. Portal-Celhay, A. Zahra, L. Kozhaya, C. Weidinger, J. Philips, D. Unutmaz, S. Feske, Ca2+ Signaling but Not Store-Operated Ca2+ Entry Is Required for the Function of Macrophages and Dendritic Cells. J Immunol 195, 1202–1217 (2015).

22. P. Li, X. Bian, C. Liu, S. Wang, M. Guo, Y. Tao, B. Huo, STIM1 and TRPV4 regulate fluid flow-induced calcium oscillation at early and late stages of osteoclast differentiation. Cell Calcium 71, 45–52 (2018).

23. H. Kajiya, F. Okamoto, T. Nemoto, K. Kimachi, K. Toh-Goto, S. Nakayana, K. Okabe, RANKL-induced TRPV2 expression regulates osteoclastogenesis via calcium oscillations. Cell Calcium 48, 260–269 (2010).

24. S.-Y. Hwang, J. W. Putney, Orai1-mediated calcium entry plays a critical role in osteoclast differentiation and function by regulating activation of the transcription factor NFATc1. FASEB J 26, 1484–1492 (2012).

25. B. Csóka, Z. H. Németh, I. Szabó, D. L. Davies, Z. V. Varga, J. Pálóczi, S. Falzoni, F. Di Virgilio, R. Muramatsu, T. Yamashita, P. Pacher, G. Haskó, Macrophage P2X4 receptors augment bacterial killing and protect against sepsis. JCI Insight 3, e99431, 99431 (2018).

26. B. N. Desai, N. Leitinger, Purinergic and calcium signaling in macrophage function and plasticity. Front Immunol 5, 580 (2014).

27. D. M. Gorman, X. X. Li, J. D. Lee, J. N. Fung, C. S. Cui, H. S. Lee, B. E. Rolfe, T. M. Woodruff, R. J. Clark, Development of Potent and Selective Agonists for Complement C5a Receptor 1 with In Vivo Activity. J Med Chem 64, 16598–16608 (2021).

28. A. Simon-Chica, A. Klesen, R. Emig, A. Chan, J. Greiner, D. Grün, A. Lother, I. Hilgendorf, E. A. Rog-Zielinska, U. Ravens, P. Kohl, F. Schneider-Warme, R. Peyronnet, Piezo1 stretch-activated channel activity differs between murine bone marrow-derived and cardiac tissue-resident macrophages. J Physiol 602, 4437–4456 (2024).

29. T. M. Link, U. Park, B. M. Vonakis, D. M. Raben, M. J. Soloski, M. J. Caterina, TRPV2 has a pivotal role in macrophage particle binding and phagocytosis. Nat Immunol 11, 232–239 (2010).

30. M. S. Schappe, K. Szteyn, M. E. Stremska, S. K. Mendu, T. K. Downs, P. V. Seegren, M. A. Mahoney, S. Dixit, J. K. Krupa, E. J. Stipes, J. S. Rogers, S. E. Adamson, N. Leitinger, B. N. Desai, Chanzyme TRPM7 Mediates the Ca2+ Influx Essential for Lipopolysaccharide-Induced Toll-Like Receptor 4 Endocytosis and Macrophage Activation. Immunity 48, 59–74.e5 (2018).

31. M. S. Schappe, M. E. Stremska, G. W. Busey, T. K. Downs, P. V. Seegren, S. K. Mendu, Z. Flegal, C. A. Doyle, E. J. Stipes, B. N. Desai, Efferocytosis requires periphagosomal Ca2+-signaling and TRPM7-mediated electrical activity. Nat Commun 13, 3230 (2022).

32. T. Murakami, J. Ockinger, J. Yu, V. Byles, A. McColl, A. M. Hofer, T. Horng, Critical role for calcium mobilization in activation of the NLRP3 inflammasome. Proc Natl Acad Sci U S A 109, 11282–11287 (2012).

33. M. Rossol, M. Pierer, N. Raulien, D. Quandt, U. Meusch, K. Rothe, K. Schubert, T. Schöneberg, M. Schaefer, U. Krügel, S. Smajilovic, H. Bräuner-Osborne, C. Baerwald, U. Wagner, Extracellular Ca2+ is a danger signal activating the NLRP3 inflammasome through G protein-coupled calcium sensing receptors. Nat Commun 3, 1329 (2012).

34. A. G. Solis, P. Bielecki, H. R. Steach, L. Sharma, C. C. D. Harman, S. Yun, N. W. Palm, M. R. de Zoete, J. N. Warnock, S. D. F. To, A. G. York, M. Mack, M. A. Schwartz, Charles S. D., Cruz, R. Jackson, R. A. Flavell, Mechanosensation of Cyclical Force by PIEZO1 is Essential for Innate Immunity. Nature 573, 69–74 (2019).

35. H. Atcha, A. Jairaman, J. R. Holt, V. S. Meli, R. R. Nagalla, P. K. Veerasubramanian, K. T. Brumm, H. E. Lim, S. Othy, M. D. Cahalan, M. M. Pathak, W. F. Liu, Mechanically activated ion channel Piezo1 modulates macrophage polarization and stiffness sensing. Nat Commun 12, 3256 (2021).

36. G. S. Baird, D. A. Zacharias, R. Y. Tsien, Circular permutation and receptor insertion within green fluorescent proteins. Proc Natl Acad Sci U S A 96, 11241–11246 (1999).

37. J. Nakai, M. Ohkura, K. Imoto, A high signal-to-noise Ca(2+) probe composed of a single green fluorescent protein. Nat Biotechnol 19, 137–141 (2001).

38. A. Miyawaki, J. Llopis, R. Heim, J. M. McCaffery, J. A. Adams, M. Ikura, R. Y. Tsien, Fluorescent indicators for Ca2+ based on green fluorescent proteins and calmodulin. Nature 388, 882–887 (1997).

39. X. R. Sun, A. Badura, D. A. Pacheco, L. A. Lynch, E. R. Schneider, M. P. Taylor, I. B. Hogue, L. W. Enquist, M. Murthy, S. S.-H. Wang, Fast GCaMPs for improved tracking of neuronal activity. Nat Commun 4, 2170 (2013).

40. T.-W. Chen, T. J. Wardill, Y. Sun, S. R. Pulver, S. L. Renninger, A. Baohan, E. R. Schreiter, R. A. Kerr, M. B. Orger, V. Jayaraman, L. L. Looger, K. Svoboda, D. S. Kim, Ultrasensitive fluorescent proteins for imaging neuronal activity. Nature 499, 295–300 (2013).

41. M. Eilert-Olsen, J. B. Hjukse, A. E. Thoren, W. Tang, R. Enger, V. Jensen, K. H. Pettersen, E. A. Nagelhus, Astroglial endfeet exhibit distinct Ca2+ signals during hypoosmotic conditions. Glia 67, 2399–2409 (2019).

42. E. Shigetomi, Y. J. Hirayama, K. Ikenaka, K. F. Tanaka, S. Koizumi, Role of Purinergic Receptor P2Y1 in Spatiotemporal Ca2+ Dynamics in Astrocytes. J Neurosci 38, 1383–1395 (2018).

43. A. J. Stevenson, G. Vanwalleghem, T. A. Stewart, N. D. Condon, B. Lloyd-Lewis, N. Marino, J. W. Putney, E. K. Scott, A. D. Ewing, F. M. Davis, Multiscale imaging of basal cell dynamics in the functionally mature mammary gland. Proceedings of the National Academy of Sciences of the United States of America 117, 26822–26832 (2020).

44. K. M. Márquez-Nogueras, E. Bovo, J. E. Neczypor, Q. Cao, A. V. Zima, I. Y. Kuo, Utilization of the genetically encoded calcium indicator Salsa6F in cardiac applications. Cell Calcium 119, 102873 (2024).

45. L. Madisen, A. R. Garner, D. Shimaoka, A. S. Chuong, N. C. Klapoetke, L. Li, A. van der Bourg, Y. Niino, L. Egolf, C. Monetti, H. Gu, M. Mills, A. Cheng, B. Tasic, T. N. Nguyen, S. M. Sunkin, A. Benucci, A. Nagy, A. Miyawaki, F. Helmchen, R. M. Empson, T. Knöpfel, E. S. Boyden, R. C. Reid, M. Carandini, H. Zeng, Transgenic mice for intersectional targeting of neural sensors and effectors with high specificity and performance. Neuron, doi: 10.1016/j.neuron.2015.02.022 (2015).

46. J. Loschko, G. J. Rieke, H. A. Schreiber, M. M. Meredith, K.-H. Yao, P. Guermonprez, M. C. Nussenzweig, Inducible targeting of cDCs and their subsets in vivo. J Immunol Methods 434, 32–38 (2016).

47. C. A. Hawley, R. Rojo, A. Raper, K. A. Sauter, Z. M. Lisowski, K. Grabert, C. C. Bain, G. M. Davis, P. A. Louwe, M. C. Ostrowski, D. A. Hume, C. Pridans, S. J. Jenkins, Csf1r-mApple Transgene Expression and Ligand Binding In Vivo Reveal Dynamics of CSF1R Expression within the Mononuclear Phagocyte System. J Immunol 200, 2209–2223 (2018).

48. B.-Z. Qian, J. Li, H. Zhang, T. Kitamura, J. Zhang, L. R. Campion, E. A. Kaiser, L. A. Snyder, J. W. Pollard, CCL2 recruits inflammatory monocytes to facilitate breast-tumour metastasis. Nature 475, 222–225 (2011).

49. R. Rojo, K. A. Sauter, L. Lefevre, D. A. Hume, C. Pridans, Maternal tamoxifen treatment expands the macrophage population of early mouse embryos. bioRxiv [Preprint] (2018). 10.1101/296749.

50. L. Deng, J.-F. Zhou, R. S. Sellers, J.-F. Li, A. V. Nguyen, Y. Wang, A. Orlofsky, Q. Liu, D. A. Hume, J. W. Pollard, L. Augenlicht, E. Y. Lin, A novel mouse model of inflammatory bowel disease links mammalian target of rapamycin-dependent hyperproliferation of colonic epithelium to inflammation-associated tumorigenesis. Am J Pathol 176, 952–967 (2010).

51. N. Taghdiri, D. M. Calcagno, Z. Fu, K. Huang, R. H. Kohler, R. Weissleder, T. P. Coleman, K. R. King, Macrophage calcium reporter mice reveal immune cell communication in vitro and in vivo. Cell Rep Methods 1, 100132 (2021).

52. J. Luo, F. Elwood, M. Britschgi, S. Villeda, H. Zhang, Z. Ding, L. Zhu, H. Alabsi, R. Getachew, R. Narasimhan, R. Wabl, N. Fainberg, M. L. James, G. Wong, J. Relton, S. S. Gambhir, J. W. Pollard, T. Wyss-Coray, Colony-stimulating factor 1 receptor (CSF1R) signaling in injured neurons facilitates protection and survival. J Exp Med 210, 157–172 (2013).

53. K. Grabert, A. Sehgal, K. M. Irvine, E. Wollscheid-Lengeling, D. D. Ozdemir, J. Stables, G. A. Luke, M. D. Ryan, A. Adamson, N. E. Humphreys, C. J. Sandrock, R. Rojo, V. A. Verkasalo, W. Mueller, P. Hohenstein, A. R. Pettit, C. Pridans, D. A. Hume, A Transgenic Line That Reports CSF1R Protein Expression Provides a Definitive Marker for the Mouse Mononuclear Phagocyte System. J Immunol 205, 3154–3166 (2020).

54. A. D. Umpierre, B. Li, K. Ayasoufi, W. L. Simon, S. Zhao, M. Xie, G. Thyen, B. Hur, J. Zheng, Y. Liang, D. B. Bosco, M. A. Maynes, Z. Wu, X. Yu, J. Sung, A. J. Johnson, Y. Li, L.-J. Wu, Microglial P2Y6 calcium signaling promotes phagocytosis and shapes neuroimmune responses in epileptogenesis. Neuron 112, 1959–1977.e10 (2024).

55. A. D. Umpierre, L. L. Bystrom, Y. Ying, Y. U. Liu, G. Worrell, L.-J. Wu, Microglial calcium signaling is attuned to neuronal activity in awake mice. Elife 9, e56502 (2020).

56. T. A. Stewart, F. M. Davis, A primary cell and organoid platform for evaluating pharmacological responses in mammary epithelial cells. ACS Pharmacology and Translational Science 3, 63–75 (2019).

57. P. Nunes, N. Demaurex, The role of calcium signaling in phagocytosis. J Leukoc Biol 88, 57–68 (2010).

58. L. P. Shutov, C. A. Warwick, X. Shi, A. Gnanasekaran, A. J. Shepherd, D. P. Mohapatra, T. M. Woodruff, J. D. Clark, Y. M. Usachev, The Complement System Component C5a Produces Thermal Hyperalgesia via Macrophage-to-Nociceptor Signaling That Requires NGF and TRPV1. J Neurosci 36, 5055–5070 (2016).

59. J. Merz, A. Nettesheim, S. von Garlen, P. Albrecht, B. S. Saller, J. Engelmann, L. Hertle, I. Schäfer, D. Dimanski, S. König, L. Karnbrock, K. Bulatova, A. Peikert, N. Hoppe, I. Hilgendorf, C. von Zur Mühlen, D. Wolf, O. Groß, C. Bode, A. Zirlik, P. Stachon, Pro- and anti-inflammatory macrophages express a sub-type specific purinergic receptor profile. Purinergic Signal 17, 481–492 (2021).

60. B. Coste, J. Mathur, M. Schmidt, T. J. Earley, S. Ranade, M. J. Petrus, A. E. Dubin, A. Patapoutian, Piezo1 and Piezo2 are essential components of distinct mechanically activated cation channels. Science 330, 55–60 (2010).

61. J. Geng, Y. Shi, J. Zhang, B. Yang, P. Wang, W. Yuan, H. Zhao, J. Li, F. Qin, L. Hong, C. Xie, X. Deng, Y. Sun, C. Wu, L. Chen, D. Zhou, TLR4 signalling via Piezo1 engages and enhances the macrophage mediated host response during bacterial infection. Nat Commun 12, 3519 (2021).

62. Y. Wang, J. Wang, J. Zhang, Y. Wang, Y. Wang, H. Kang, W. Zhao, W. Bai, N. Miao, J. Wang, Stiffness sensing via Piezo1 enhances macrophage efferocytosis and promotes the resolution of liver fibrosis. Sci Adv 10, eadj3289 (2024).

63. S. Pourteymour, J. Fan, R. K. Majhi, S. Guo, X. Sun, Z. Huang, Y. Liu, H. Winter, A. Bäcklund, N.-T. Skenteris, E. Chernogubova, O. Werngren, Z. Li, J. Skogsberg, Y. Li, L. Matic, U. Hedin, L. Maegdefessel, E. Ehrenborg, Y. Tian, H. Jin, PIEZO1 targeting in macrophages boosts phagocytic activity and foam cell apoptosis in atherosclerosis. Cell Mol Life Sci 81, 331 (2024).

64. S. M. Swain, R. A. Liddle, Piezo1 acts upstream of TRPV4 to induce pathological changes in endothelial cells due to shear stress. J Biol Chem 296, 100171 (2021).

65. A. J. Hyman, S. Tumova, D. J. Beech, Piezo1 Channels in Vascular Development and the Sensing of Shear Stress. Current Topics in Membranes, doi: 10.1016/bs.ctm.2016.11.001 (2017).

66. S. S. Ranade, Z. Qiu, S.-H. S. H. Woo, S. S. Hur, S. E. Murthy, S. M. Cahalan, J. Xu, J. Mathur, M. Bandell, B. Coste, Y. S. J. Y.-S. J. Li, S. Chien, A. Patapoutian, Piezo1, a mechanically activated ion channel, is required for vascular development in mice. 111, 10347–10352 (2014).

67. J. Li, B. Hou, S. Tumova, K. Muraki, A. Bruns, M. J. Ludlow, A. Sedo, A. J. Hyman, L. McKeown, R. S. Young, N. Y. Yuldasheva, Y. Majeed, L. A. Wilson, B. Rode, M. A. Bailey, H. R. Kim, Z. Fu, D. A. L. Carter, J. Bilton, H. Imrie, P. Ajuh, T. N. Dear, R. M. Cubbon, M. T. Kearney, K. R. Prasad, P. C. Evans, J. F. X. Ainscough, D. J. Beech, Piezo1 integration of vascular architecture with physiological force. Nature 515, 279–282 (2014).

68. J. Richardson, A. Kotevski, K. Poole, From stretch to deflection: the importance of context in the activation of mammalian, mechanically activated ion channels. FEBS J 289, 4447–4469 (2022).

69. S. Ma, A. E. Dubin, Y. Zhang, S. A. R. Mousavi, Y. Wang, A. M. Coombs, M. Loud, I. Andolfo, A. Patapoutian, A role of PIEZO1 in iron metabolism in mice and humans. Cell 184, 969–982.e13 (2021).

70. Z. Zhou, X. Ma, Y. Lin, D. Cheng, N. Bavi, G. A. Secker, J. V. Li, V. Janbandhu, D. L. Sutton, H. S. Scott, M. Yao, R. P. Harvey, N. L. Harvey, B. Corry, Y. Zhang, C. D. Cox, MyoD-family inhibitor proteins act as auxiliary subunits of Piezo channels. Science 381, 799–804 (2023).

71. M. R. Servin-Vences, M. Moroni, G. R. Lewin, K. Poole, Direct measurement of TRPV4 and PIEZO1 activity reveals multiple mechanotransduction pathways in chondrocytes. eLife 6, e21074 (2017).

72. S. Shrestha, J. Richardson, K. Poole, Analysing Mechanically Evoked Currents at Cell-Substrate Junctions. Methods Mol Biol 2600, 155–167 (2023).

73. K. Poole, R. Herget, L. Lapatsina, H.-D. Ngo, G. R. Lewin, Tuning Piezo ion channels to detect molecular-scale movements relevant for fine touch. Nat Commun 5, 3520 (2014).

74. N. Bavi, J. Richardson, C. Heu, B. Martinac, K. Poole, PIEZO1-Mediated Currents Are Modulated by Substrate Mechanics. ACS Nano 13, 13545–13559 (2019).

75. J. W. Pollard, L. Hennighausen, Colony stimulating factor 1 is required for mammary gland development during pregnancy. Proceedings of the National Academy of Sciences 91, 9312–9316 (1994).

76. S. Shindo, S. Nakamura, M. Rawas-Qalaji, A. Heidari, M. R. Pastore, M. Okamoto, M. Suzuki, M. Salinas, D. Minond, A. Bontempo, M. Cayabyab, Y. Yang, J. L. Crane, M. Hernandez, S. Vardar, P. Hardigan, X. Han, S. Kaltman, T. Kawai, Mechanosensing by Piezo1 regulates osteoclast differentiation via PP2A-Akt axis in periodontitis. bioRxiv [Preprint] (2024). 10.1101/2024.09.04.611049.

77. W. Sun, S. Chi, Y. Li, S. Ling, Y. Tan, Y. Xu, F. Jiang, J. Li, C. Liu, G. Zhong, D. Cao, X. Jin, D. Zhao, X. Gao, Z. Liu, B. Xiao, Y. Li, The mechanosensitive Piezo1 channel is required for bone formation. Elife 8, e47454 (2019).

78. K. A. Sauter, C. Pridans, A. Sehgal, Y. T. Tsai, B. M. Bradford, S. Raza, L. Moffat, D. J. Gow, P. M. Beard, N. A. Mabbott, L. B. Smith, D. A. Hume, Pleiotropic effects of extended blockade of CSF1R signaling in adult mice. J Leukoc Biol 96, 265–274 (2014).

79. S. Uderhardt, A. J. Martins, J. S. Tsang, T. Lämmermann, R. N. Germain, Resident Macrophages Cloak Tissue Microlesions to Prevent Neutrophil-Driven Inflammatory Damage. Cell 177, 541–555.e17 (2019).

80. J. R. Holt, W.-Z. Zeng, E. L. Evans, S.-H. Woo, S. Ma, H. Abuwarda, M. Loud, A. Patapoutian, M. M. Pathak, Spatiotemporal dynamics of PIEZO1 localization controls keratinocyte migration during wound healing. Elife 10, e65415 (2021).

81. G. Jacquemin, M. Benavente-Diaz, S. Djaber, A. Bore, V. Dangles-Marie, D. Surdez, S. Tajbakhsh, S. Fre, B. Lloyd-Lewis, Longitudinal high-resolution imaging through a flexible intravital imaging window. Science Advances 7, eabg7763 (2021).

82. L. Mourao, M. Ciwinska, J. van Rheenen, C. L. G. J. Scheele, Longitudinal Intravital Microscopy Using a Mammary Imaging Window with Replaceable Lid. J Vis Exp, doi: 10.3791/63326 (2022).

83. D. A. Hume, The many alternative faces of macrophage activation. Frontiers in Immunology 6 (2015).

84. F. M. Davis, I. Azimi, R. A. Faville, A. A. Peters, K. Jalink, J. W. Putney, G. J. Goodhill, E. W. Thompson, S. J. Roberts-Thomson, G. R. Monteith, Induction of epithelial-mesenchymal transition (EMT) in breast cancer cells is calcium signal dependent. Oncogene 33, 2307–2316 (2014).

85. T. B.-D. McEwan, R. A. Sophocleous, P. Cuthbertson, K. J. Mansfield, M. L. Sanderson-Smith, R. Sluyter, Autocrine regulation of wound healing by ATP release and P2Y2 receptor activation. Life Sci 283, 119850 (2021).

86. C. Wetzel, S. Pifferi, C. Picci, C. Gök, D. Hoffmann, K. K. Bali, A. Lampe, L. Lapatsina, R. Fleischer, E. S. J. Smith, V. Bégay, M. Moroni, L. Estebanez, J. Kühnemund, J. Walcher, E. Specker, M. Neuenschwander, J. P. von Kries, V. Haucke, R. Kuner, J. F. A. Poulet, J. Schmoranzer, K. Poole, G. R. Lewin, Small-molecule inhibition of STOML3 oligomerization reverses pathological mechanical hypersensitivity. Nat Neurosci 20, 209–218 (2017).

87. L. Wang, X. You, S. Lotinun, L. Zhang, N. Wu, W. Zou, Mechanical sensing protein PIEZO1 regulates bone homeostasis via osteoblast-osteoclast crosstalk. Nat Commun 11, 282 (2020).

88. J. Schindelin, I. Arganda-Carreras, E. Frise, V. Kaynig, M. Longair, T. Pietzsch, S. Preibisch, C. Rueden, S. Saalfeld, B. Schmid, J.-Y. J.-Y. Tinevez, D. J. White, V. Hartenstein, K. Eliceiri, P. Tomancak, A. Cardona, K. Liceiri, P. Tomancak, C. A., Fiji: an open source platform for biological image analysis. Nature Methods 9, 676–682 (2012).

89. C. Stringer, M. Pachitariu, Cellpose3: one-click image restoration for improved cellular segmentation. Nat Methods 22, 592–599 (2025).

90. T. A. Stewart, F. M. Davis, Got milk? Identifying and characterizing lactation defects in genetically-engineered mouse models. Journal of Mammary Gland Biology and Neoplasia (2020).

91. B. Lloyd-Lewis, F. M. Davis, O. B. Harris, J. R. Hitchcock, F. C. Lourenco, M. Pasche, C. J. Watson, Imaging the mammary gland and mammary tumours in 3D: Optical tissue clearing and immunofluorescence methods. Breast Cancer Research 18 (2016).

92. M.-T. Ke, S. Fujimoto, T. Imai, SeeDB: a simple and morphology-preserving optical clearing agent for neuronal circuit reconstruction. Nature neuroscience 16, 1154–61 (2013).

93. M. Linkert, C. T. Rueden, C. Allan, J. M. Burel, W. Moore, A. Patterson, B. Loranger, J. Moore, C. Neves, D. MacDonald, A. Tarkowska, C. Sticco, E. Hill, M. Rossner, K. W. Eliceiri, J. R. Swedlow, Metadata matters: access to image data in the real world. Journal of Cell Biology 189, 777–782 (2010).

94. K.-U. Wagner, C. a Boulanger, M. D. Henry, M. Sgagias, L. Hennighausen, G. H. Smith, An adjunct mammary epithelial cell population in parous females: its role in functional adaptation and tissue renewal. Development (Cambridge, England) 129, 1377–86 (2002).

95. T. A. Stewart, K. Hughes, A. J. Stevenson, N. Marino, A. L. Ju, M. Morehead, F. M. Davis, Mammary mechanobiology - investigating roles for mechanically activated ion channels in lactation and involution. Journal of cell science 134 (2021).

96. E. A. Susaki, K. Tainaka, D. Perrin, H. Yukinaga, A. Kuno, H. R. Ueda, Advanced CUBIC protocols for whole-brain and whole-body clearing and imaging. Nat Protoc 10, 1709–1727 (2015).

97. K. M. Suchanek, F. J. May, J. A. Robinson, W. J. Lee, N. A. Holman, G. R. Monteith, S. J. Roberts-Thomson, Peroxisome proliferator-activated receptor alpha in the human breast cancer cell lines MCF-7 and MDA-MB-231. Molecular carcinogenesis 34, 165–171 (2002).

98. M. Folacci, S. B. Chalmers, F. M. Davis, “Methods for Imaging Intracellular Calcium Signals in the Mouse Mammary Epithelium in Two and Three Dimensions” in Calcium Signaling: Methods and Protocols, C. M. Gorvin, Ed. (Springer US, New York, NY, 2025; 10.1007/978-1-0716-4164-4_15), pp. 195–212.

99. F. M. Davis, A. Janoshazi, K. S. Janardhan, N. Steinckwich, D. M. D’Agostin, J. G. Petranka, P. N. Desai, S. J. Roberts-Thomson, G. S. Bird, D. K. Tucker, S. E. Fenton, S. Feske, G. R. Monteith, J. W. Putney, Essential role of Orai1 store-operated calcium channels in lactation. Proceedings of the National Academy of Sciences 112, 5827–5832 (2015).

100. S. Sianati, A. Kurumlian, E. Bailey, K. Poole, Analysis of Mechanically Activated Ion Channels at the Cell-Substrate Interface: Combining Pillar Arrays and Whole-Cell Patch-Clamp. Front Bioeng Biotechnol 7, 47 (2019).

101. M. B. Brent, J. S. Thomsen, A. Brüel, Short-term glucocorticoid excess blunts abaloparatide-induced increase in femoral bone mass and strength in mice. Sci Rep 11, 12258 (2021).

